# HiExM Enables Scalable Mapping of Organelle Morphology and Spatial Heterogeneity

**DOI:** 10.64898/2026.07.12.738053

**Authors:** John H. Day, John Farrell, Dandan Yang, Fabricio N. Neira, Elizabeth A. Allen, Ann M. Byrne, Nina C. Leksa, Katherine W. Klinger, Giovanni de Nola, Mushriq Al-Jazrawe, Laurie A. Boyer

## Abstract

Quantitative image analysis of subcellular organization requires sufficient spatial resolution to resolve individual organelles and sample size to capture heterogeneity both within cells and between cells. Existing imaging approaches often force a tradeoff between spatial resolution and throughput, limiting the ability to measure organelle-level phenotypes across cell populations. Here, we establish high-throughputs expansion microscopy (HiExM) as a scalable pipeline for single-organelle analysis. As a benchmark, we focus on mapping late endosomes and lysosomes (LELs), a heterogeneous organelle class whose small size, dense intracellular distribution, and functional diversity make it difficult to quantify accurately using conventional light microscopy. HiExM increases effective spatial resolution while preserving compatibility with large-scale image acquisition, enabling robust segmentation and quantitative profiling of individual LELs across large cell populations. Using this pipeline, we identified differences in intracellular trafficking behavior among anti-transferrin receptor antibodies that could not be captured by conventional colocalization analysis alone. We further integrate spatial and morphological features with learned image-based representations that can define relationships between LEL morphology and subcellular position as well as how these relationships respond to perturbations. Together, our work establishes HiExM as a generalizable platform for scalable single-organelle profiling, enabling an analytical framework for quantifying discrete organelles across cells and conditions.

## Introduction

Quantitative analysis of subcellular organization requires imaging approaches that can resolve individual structures while supporting measurements across large cell populations^1^. A major technical barrier is that many organelles exist at spatial scales near or below the diffraction limit of light microscopy, limiting the ability to distinguish closely spaced objects and the throughput required for statistically robust analysis^2^. For example, conventional fluorescence microscopy often produces merged or ambiguous signals, preventing accurate identification and segmentation of individual organelles required for downstream computational approaches^3^.

Modern computational approaches for image-based cell biology, including object-based feature extraction, spatial statistics, and machine learning, require accurate segmentation of individual organelles^4,5^. Without reliable segmentation, analyses are restricted to pixel-level or bulk measurements (e.g., global intensity or colocalization), which obscure measurements of organelle-level variability and reduce sensitivity to detecting phenotypic differences. While electron microscopy provides nanometer-scale resolution (<10 nm) and resolves organelle ultrastructure, this approach is less readily compatible with scalable, multiplexed, protein-specific measurement^6,7^. Super-resolution fluorescence microscopy improves effective resolution to ∼20–50 nm, but can be limited by acquisition speed, field of view, or throughput, restricting its application to small-scale studies^3^. Consequently, high resolution analysis of discrete organelles across many cells in a high-throughput context remains a challenge.

Expansion microscopy (ExM) approaches address these key bottlenecks by physically enlarging biological specimens, enabling effective resolutions 4-fold higher than standard diffraction-limited microscopes^8,9^. Importantly, ExM preserves fluorescence labeling and molecular specificity, allowing multi-channel imaging of subcellular structures. Moreover, advances in high-throughput expansion microscopy (HiExM) enable parallelized sample processing in multi-well formats compatible with automated imaging platforms^10^.

Here, we developed an integrated framework that combines HiExM and robust single-organelle segmentation of lysosomes with a suite of computational analyses to enable quantitative, organelle-resolved imaging at scale. As a benchmark, the lysosomal compartment comprises heterogeneous organelles with various cellular roles that range from ∼100 nm to >1 µm in diameter and are frequently separated by distances below ∼200 nm^6,11,12^. While lysosome-associated membrane protein 1 (LAMP1) is well-suited for global labeling of both late endosomes and lysosomes (LELs), its broad distribution emphasizes the need for increased spatial resolution to distinguish organelle features^13^. We demonstrate that HiExM can resolve individual organelles in densely packed cellular environments by enabling accurate segmentation. Our object-level analysis reveals differences in cellular localization over time when comparing three anti-transferrin receptor (anti-TfR) antibody variants. We also integrate segmentation outputs with machine learning, including interpretable feature-based measurements and unsupervised representation learning. This approach enables extraction of quantitative descriptors of organelle size, spatial positioning, and high-dimensional morphological variation across large cell populations. We also demonstrate that our pipeline identifies perturbation-induced morphological phenotypes. Together, this work establishes a deployable strategy for integrating nanoscale imaging with quantitative analysis of single organelles, which could advance our understanding of cell biology and enhance drug screening applications.

## Results

### HiExM Enables Mapping of Single LAMP1 Organelles

Standard fluorescence microscopy is fundamentally limited to ∼200 nm resolution, but often provides much coarser resolution depending on the microscope, limiting observations of subcellular structures^2^. Given that LELs generally range from 100-1000+ nm in size and are often densely organized inside cells, we applied HiExM to map individual LELs within and across cells. Using LAMP1 as a broad marker of these organelles, we show expansion in hCMECs results in the expected ∼4-fold expansion and similar levels of gel distortion compared to prior expansion microscopy reports (Supplementary Fig. 1)^8–10^. Comparison between the pre-and post-expansion images showed a dramatic improvement in the resolution of LAMP1+ organelles (Fig. 1A-C). While the increase in lateral resolution is important in resolving individual organelles, the 4-fold increase in axial resolution was critical to disambiguate individual organelles that are vertically stacked within cells (Fig. 1B-C). This improved resolution yields discrete LAMP1 compartments that are identifiable by eye. Using post-expansion images, we manually generated ground truth segmentations of LELs and trained a StarDist model for segmentation (Fig. 1D, Supplementary Fig. 2)^14–16^. StarDist was chosen because it is designed for robust segmentation of objects with relatively regular, somewhat spherical, star-convex shapes consistent with the organelles analyzed in this work.

**Fig. 1:**
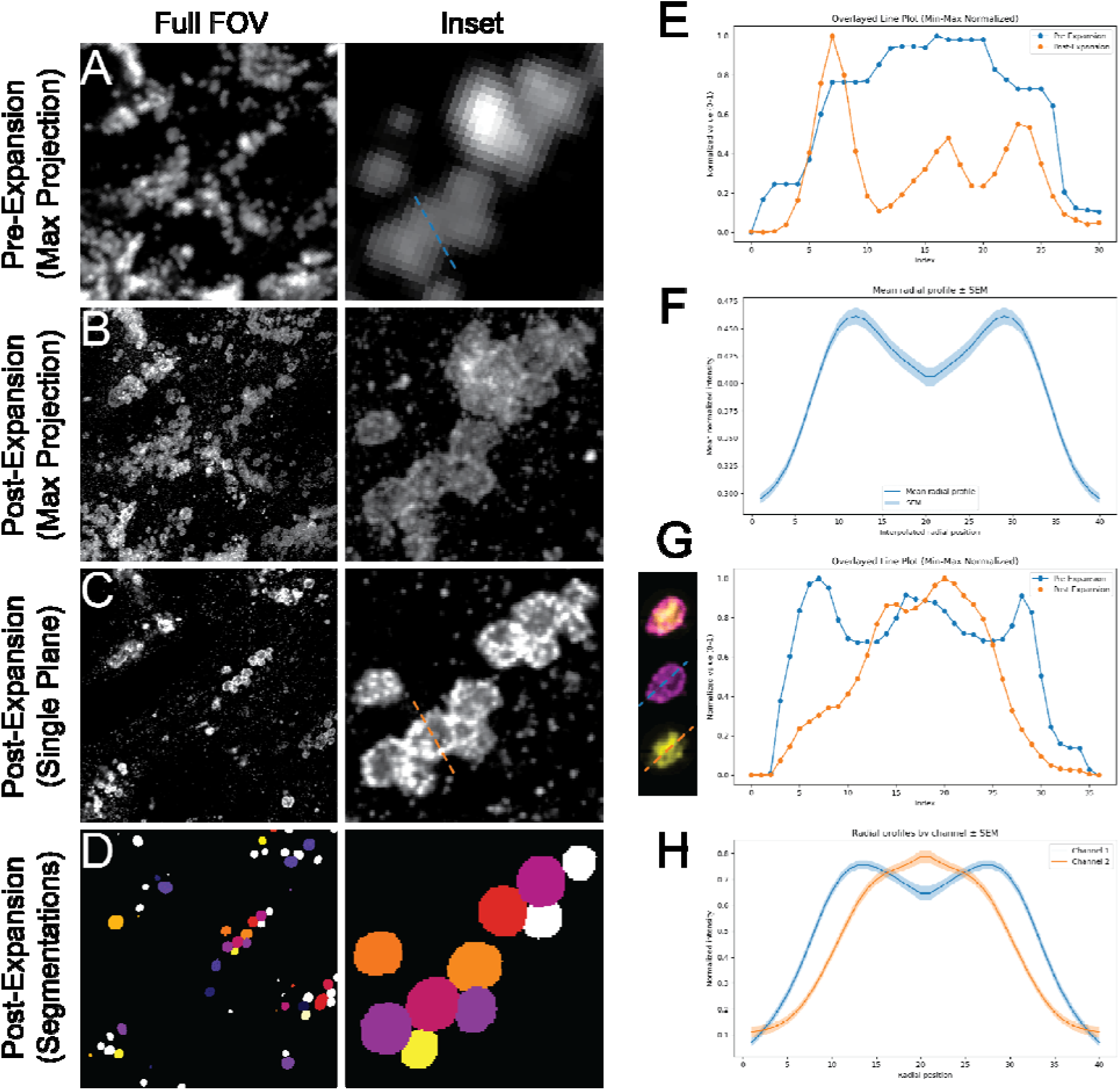
High-throughput Expansion microscopy (HiExM) enables LEL segmentation and single-organelle analytics. A) Representative pre-expansion LAMP1 image maximum projection. B) The same field-of-view as in A, taken post-expansion, shown as a maximum projection. C) The same field of view shown as a single optical section from the z-stack. D) Application of the StarDist segmentation model on this same optical section, post-expansion. E) Line plot from dotted lines shown in the insets for A and C. F) Mean radial intensity profile of 241 LAMP1-positive organelles. Distances were normalized to allow averaging across organelles of different sizes, and the profile is displayed symmetrically such that the plot edges correspond to the organelle boundary and the plot center corresponds to the center of the organelle. G) (Left) Example image of a single segmented LAMP1-positive (magenta) and LAMP3-positive organelle (yellow). (right) Line plot showing the signal intensities of LAMP1 and LAMP3 for the example organelle along the dotted lines in the image. H) Mean radial intensity profiles of LAMP1 and LAMP3 measured across segmented LELs, with distances normalized to organelle size as in F.

A defining feature of LELs is the presence of a distinct lumen. In most cases, observing this feature requires higher image resolution than standard confocal microscopy (Fig. 1E, Supplementary Figure 3). To demonstrate resolution, we used a ring-based analysis at the center plane of each segmented organelle to map the mean signal intensity as a function of distance from the periphery of the masked organelle. By normalizing the scale of the distance metric, we aggregated data from 241 segmented LAMP1 organelles of varying sizes. This analysis reveals a consistently distinguishable lumen in organelles across the dataset (Fig. 1F). Next, we evaluated the distribution of CD63 (also known as LAMP3), a lysosome-associated protein that is known to colocalize with LAMP1, within LELs^13,17^. Electron microscopy studies have shown that LAMP3 localizes predominantly on the membranes of intralumenal vesicles^18^. To determine whether this organization could be captured by HiExM, we stained cells with LAMP1 and LAMP3 antibodies, expanded the samples, and analyzed signal distributions in the LAMP1 segmentations. As shown in a representative example, we observe LAMP3 signal concentrated inside the lumen while LAMP1 signal is highest at the organelle membrane (Fig. 1G). When aggregated across a dataset of 241 organelles, this pattern remains consistent (Fig. 1H), indicating that HiExM can resolve sub-organelle organization within the lysosomal compartment.

### Single-Organelle Segmentations Enable Precise Mapping of Antibody uptake in the LEL Compartment

Having established our approach for resolving LAMP1 organelles, we next applied this framework to analyze cellular uptake of anti-TfR antibodies. Anti-TfR antibodies have recently been used as drug vehicles because of their ability to ferry tethered cargo into the cell via clathrin-mediated endocytosis^19^. Characterizing the intracellular distribution of these antibodies over time is challenging because endocytic organelles including LELs exist at length scales that are difficult to resolve with conventional imaging approaches^6^. To resolve antibody localization within the LELs, hCMECs were treated with anti-TfR antibodies to compare the localization of a bivalent anti-TfR antibody (AbB) and monovalent anti-TfR antibody (AbM)^20^. Briefly, cells were incubated with antibody and fixed at 30-minute intervals over five hours followed by staining and application of the HiExM protocol. Following expansion, we designed a complementary set of analyses to test three key performance criteria: sensitivity to detect differences in antibody localization, spatial precision to assign antibody signal to segmented LEL volumes, and single-organelle resolution to distinguish whether cargo accumulates broadly across the LEL compartment or within discrete organelle subsets.

We first performed a signal-based colocalization analysis on loosely masked whole image fields-of-view using the Pearson-correlation coefficient (PCC). Masks were generated for LAMP1 signal using adaptive thresholding to isolate the general volume in and around the LAMP1 signal in images. PCC quantifies the degree to which voxel intensities in two channels covary across an image, yielding a measure of the similarity in spatial distribution between them. Using standard confocal microscopy, we could not detect a difference in anti-TfR antibody colocalization with LAMP1 between AbB and AbM (Fig. 2A,B). Conversely in HiExM, AbB shows greater colocalization with LAMP1 than AbM as early as 90 minutes by PCC (Fig. 2C,D). Additionally, the post-expansion images show that neither antibody colocalizes (PCC<0) with LAMP1 before 90 minutes, consistent with the timing of Ab trafficking to the lysosome^21^. Notably, we found that a fusion protein of AbM and alglucosidase alfa (AbM-GAA)^20^ strongly colocalized with LAMP1 organelles compared to either cargo-free antibody (Supplementary Fig. 4), indicating the high sensitivity of our assay to detect localization differences.

**Fig. 2:**
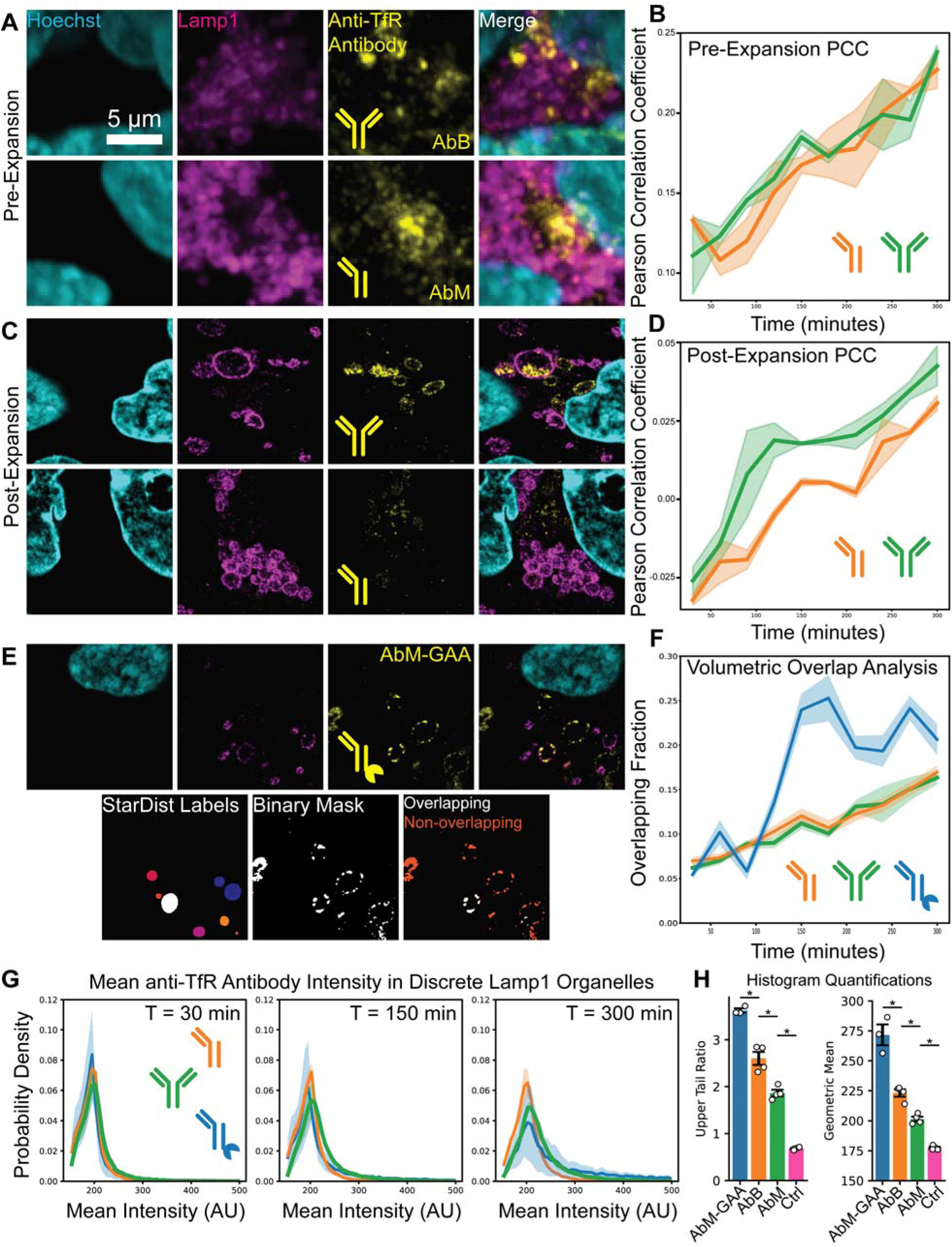
HiExM Enables High-resolution Colocalization of LAMP1 and Anti-TFR Antibody Signals. A) Standard confocal microscopy yields images with ambiguous signal overlap. B) Standard image resolution is insufficient to ascertain a difference in colocalization between monovalent and bivalent anti-TfR antibodies over time. C) The added resolution from HiExM yields fine detail for single-organelle identification and signal overlap evaluation. D) Differences in colocalization with LAMP1 over time are observed between the monovalent and bivalent anti-TfR antibodies. E) Example of volumetric overlap analysis. LELs are segmented with StarDist and a binary mask is applied to the drug channel with an absolute threshold. The overlap between LAMP1 StarDist segmentations and anti-TfR-positive voxels is then evaluated. F) Colocalization based on volumetric overlap between LAMP1 segmentations and anti-TfR binary masks shows no difference between cargo-free anti-TfR antibodies, while AbM-GAA fusion antibodies show greater volumetric overlap. G) Object-based intensity analysis identifies differences in colocalization among antibodies. Histograms comparing the anti-TfR antibodies tested at three different timepoints are shown. Each curve shows the mean distribution of LAMP1 organelles in terms of their mean intensity in the drug channel. H) Key histogram metrics that differentiate anti-TfR antibody types. The upper-tail Ratio (left) represents the difference between the 95^th^ percentile value and the median value divided by the interquartile range. This value indicates how far the top end of the distribution is compared to the median. The geometric mean was also calculated. Error bars and shaded regions, SEM of the means from four replicate wells (∼10 fields-of-view per well) taken across two experiments. *Significant (p<0.05) by Welch’s two-sample t test, tested on the mean values from three-four independent replicates (wells) across two experiments.

We next used segmentation to analyze colocalization in organelle volumes. LEL segmentations enabled us to generate estimates of the relative proportion of anti-TfR antibody signal within or excluded from LAMP1 volumes. For this analysis, an absolute threshold was applied to antibody images and the number of antibody-positive voxels overlapping segmented LEL volumes was compared with the total number of antibody-positive voxels to estimate the proportion of antibody signal associated with LELs (Fig. 2E). It is important to note that this value does not take into account the signal intensity in the drug voxels. Interestingly, while this volumetric analysis showed higher anti-TfR antibody-LAMP1 overlap with AbM-GAA compared to either cargo-free antibody, little to no difference was observed between AbM and AbB (Fig. 2F). Contrasted with the PCC result, these data suggest that the AbM and AbB antibodies occupy LAMP1-associated volumes to a similar extent, but that the bivalent antibody exhibits stronger LAMP1-colocalized signal intensity, consistent with earlier localization to LELs during the time course as shown above.

We next calculated mean anti-TfR antibody signal intensity on a per-LEL basis and compared the anti-TfR antibodies at different timepoints. Using these data, we plotted the distributions of single LELs based on mean anti-TfR antibody signal intensity (Fig. 2G). These results reveal two key observations that differentiate the intracellular behaviors of AbM-GAA and AbB, both of which colocalize with LAMP1 more prominently than AbM. First, the histogram of the bivalent antibody exhibits an up-shift in the geometric mean signal intensity value, suggesting a global increase in AbB localization across LELs over time. In contrast, the histogram of AbM-GAA exhibits a lengthened tail, indicating that AbM-GAA resides in a smaller population of LELs (Fig. 2H). Notably, we also show that AbM-GAA could rescue the lysosomal hypertrophy phenotype in cardiomyocytes from Pompe patient-derived induced pluripotent stem cells (iPSCs) (see Supplementary Methods). Our results show that, while both AbM-GAA and GAA alone rescue the lysosomal hypertrophy phenotype and improve sarcomere organization in Pompe cardiomyocytes, the non-cardiomyocytes in these cultures required AbM-GAA for phenotypic rescue (Supplementary Fig. 5). Collectively, these analyses demonstrate that HiExM meets our key benchmarks for quantitative antibody-localization by resolving whether anti-TfR antibodies reach the LEL compartment and reveal how different antibodies could shape their distribution across LELs. Thus, the ability to measure these attributes at scale could improve the rational design of new antibody-based drugs optimized for specific cellular outcomes^22^.

### Single-Organelle Mapping Reveals Relationships Between LEL Morphology and Spatial Positioning

Because studies suggest that collective features including organelle morphology, spatial clustering and cellular localization are critical for assessing lysosomal diversity^12,23^, we next developed a general framework for the systematic classification and characterization of these organelles. To first validate organelle heterogeneity using an orthogonal method, we used Tokuyasu-Correlative Light/Electron Microscopy (TO-CLEM) to qualitatively evaluate LAMP1-positive LELs in hCMECs. As expected, TO-CLEM images show LAMP1 signal overlaps with multivesicular bodies and multilamellar lysosomes (Supplementary Fig. 6; Supplementary Methods)^12,23^. Thus, we tested whether HiExM analysis could provide richer information of LEL diversity.

We focused our analysis on a set of five features including volume, nearest distance to the nucleus, nearest distance to the cell membrane, number of proximal LELs and number of branches in a minimal graph (see Methods) for each LEL using HiExM images. These features were chosen to capture organelle size, intracellular positioning, and local spatial organization. Each of these features has been associated with differences in lysosome function, trafficking behavior, or interaction with neighboring compartments^24–27^. To delineate cell boundaries, because standard cell-surface stains (e.g., wheat-germ agglutinin) were challenging with expansion microscopy, we performed transient transfection of eGFP and staining to label the cytoplasmic compartment prior to expansion and used the resulting fluorescence signal to mask single-cell volumes (Fig. 3A). This approach allowed us to map the relative positions and diversity of LELs across 89 hCMEC cells (Fig. 3B). Based on our measured features of all LELs, these organelles fell into 14 clusters (Supplementary Fig. 7). Leiden clustering was used for this analysis because we had no a priori expectation that our data would form clusters in feature space with simple geometric structure^28^. Because Leiden identifies communities based on local neighborhood connectivity, it is well suited to irregular manifolds such as those often observed in biological data. Our data show that LELs from any given cell are distributed across most of the clusters meaning that all cells contain LELs spanning a wide range of sizes, relative spatial positions, and local clustering regimes (Supplementary Fig. 8). This broad intracellular distribution of LEL phenotypes indicates that heterogeneity is an intrinsic feature of the LEL population in hCMECs. Notably, we find that clusters with high organelle volume correlate with higher numbers of connected objects. Conversely, the clusters characterized by high distance from the nucleus also represent the smallest and least connected LELs. These patterns are consistent with observations that larger LELs exhibit spatial clustering in cells and peripheral LELs are generally smaller and sparser^21,26^. Motivated by this result, we next developed a generalized approach to probe the relationship between LEL cellular positioning and morphology.

**Fig. 3:**
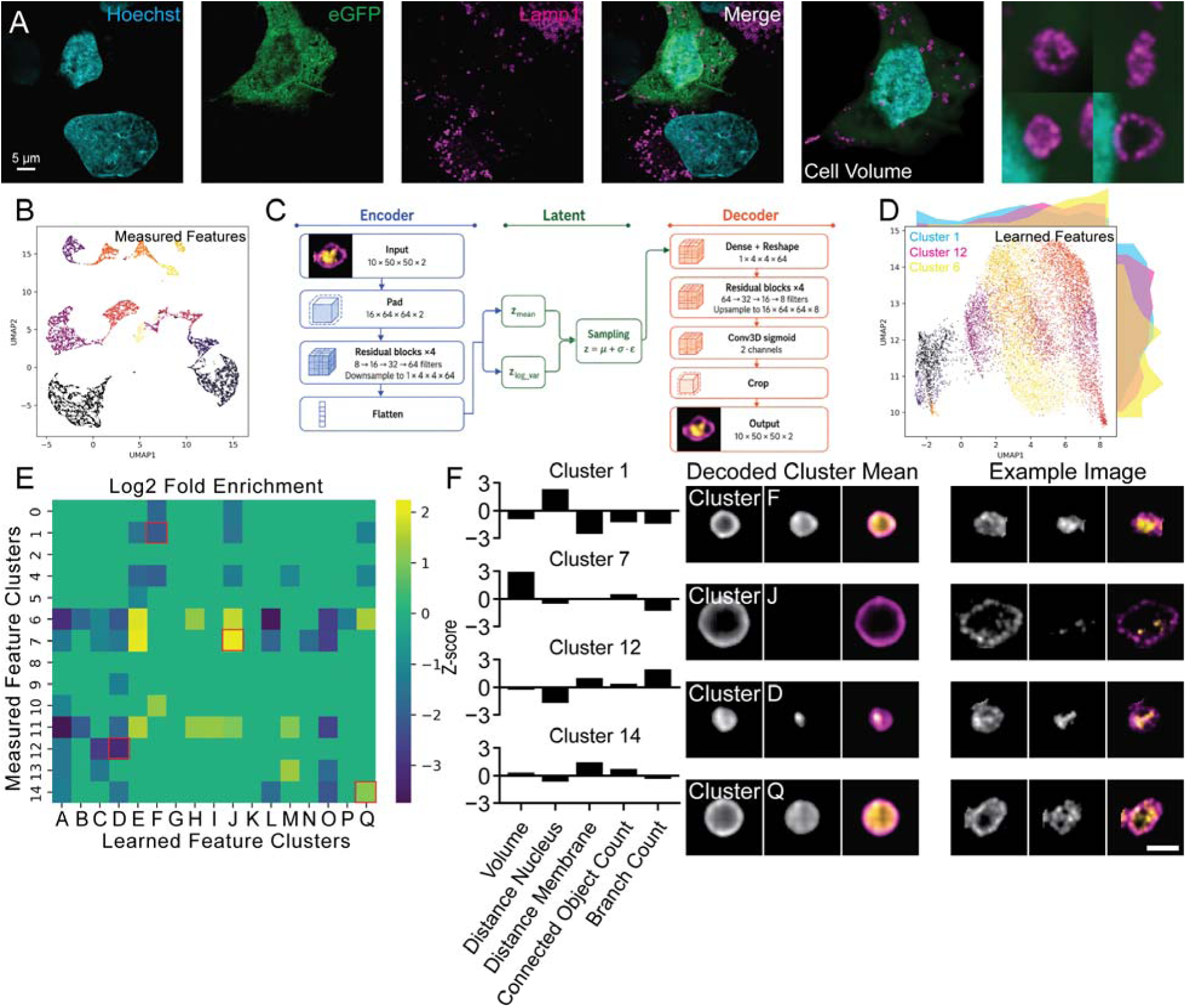
LEL Measured and Learned Features Reveal Spatial and Morphological Profiles of LELs in Single Cells. A) Example single-cell image stained for Hoechst (cyan), eGFP (green), and LAMP1 (Magenta). Insets on the right show example single-plane images of individual LAMP1 organelles in this cell. B) UMAP of measured features across 13271 LELs from untreated cells (eight wells from two independent experiments, ∼10 fields-of-view per well capturing one cell each, 89 cells total). C) Abridged architecture of the convolutional neural network based variational autoencoder used to vectorize segmented LEL images. A real example input and decoded result is shown. A schematic of the model architecture can be found in Supplementary Fig. 9. D) UMAP of learned features across LELs from untreated cells. Histograms on the edges of the UMAP represent the distributions of clusters 1, 6, and 12 from the measured features in the learned feature UMAP space along UMAP1 and UMAP2. E) Log2-fold enrichment plot comparing measured and learned feature clusters. For each combination of a measured and learned feature cluster, the number of objects that fall into both clusters was compared to the expected number of objects that would fall into both clusters given random chance. Only clusters with greater than 2-fold enrichment or depletion are shown for simplicity. F) Example pairings of measured (left) and Learned (right) feature cluster identities. The measured feature clusters are identified by a characteristic set of z-score values for the 5 measured features. The learned feature clusters are identified by either taking the mean feature vector for all of the vectors in that cluster and decoding it (left) or taking a representative example organelle from that cluster (right). Scale bar: 1 µm.

To quantify LEL morphology, we built a variational autoencoder based on a previously published architecture (Supplementary Fig. 9)^1,29^. Briefly, this model was trained on 100,000 individual segmented image volumes of LELs stained for LAMP1 and LAMP3 (Fig. 1G) to provide high-quality and information-rich image data for machine learning. The model was trained to minimize the mean-squared error between the original and reconstructed images, resulting in a 256-dimensional latent space that encodes minimal representations of LEL images while retaining their key features (Fig. 3C). After model training, validation, and testing, we used the encoder-half of the model to vectorize LEL images, providing an orthogonal data modality to probe LEL morphology independently of the measured features. As an initial validation of the learned feature space, we assessed the distributions of LAMP1 and LAMP3 signal intensities (Supplementary Fig. 10). The structured patterning of these signal intensity values supports the idea that the learned representation encodes informative features of the organelles. As above, we clustered LEL learned-feature vectors into 16 clusters, each representing a morphologically distinct subset of organelles (Fig. 3D, Supplementary Fig. 11).

Because cellular positioning and organelle morphology have both been independently linked with LEL functional state^24,25^, we reasoned that these features could be related to one another. To this end, plotting the distribution of three measured-feature clusters on UMAP1 and UMAP2 in our reduced learned-feature space showed that these clusters exhibit distinct and non-uniform distributions (Fig. 3D). This pattern is consistent with the notion that organelle morphology and spatial distribution characteristics are related. To further examine the relationship between LEL morphology and spatial distribution, we identified combinations of clusters that were enriched or depleted in LEL abundance relative to the expectation under a null model of random association (Fig. 3E). We observed multiple cluster combinations with greater than twofold enrichment or depletion, indicating that certain spatially defined LEL states are preferentially associated with specific learned morphological classes at baseline (Fig. 3F). As an internal validation of this approach, measured-feature cluster 7 characterized by large-volume LELs, was strongly enriched for learned-feature cluster J that likewise corresponds to relatively large organelles. Together, these results indicate that the learned and measured features capture related aspects of the LEL state.

### Learned LEL Features Reveal Differences Between Distinct Lysosomal Perturbations

To benchmark the ability of HiExM to detect changes in the above LEL features, we next tested how perturbing lysosome function through multiple distinct mechanisms would impact our learned LEL features. We treated hCMECs with apilimod, bafilomycin, and chloroquine compared to control treatment with DMSO (Fig. 4A, Supplementary Table 2). Each of these compounds impacts lysosomal function through a unique mechanism of action^30,31^. After compound treatment, we performed HiExM and analyzed 73,890 LELs across 332 cells by generating clusters based on measured and learned features as before (Fig. 4B,C). To compare perturbations, we calculated the fold change in LEL abundance relative to control for each cluster across the three compounds in both the measured and learned feature sets (Fig. 4D,E). Interestingly, in the measured-feature space, no cluster exhibited more than a twofold difference between any pair of compounds. Conversely, the learned-feature space shows several clusters with greater than twofold differential enrichment/depletion between conditions, indicating stronger compound-specific differences in learned LEL morphology. This result suggests that, although these compounds produce broadly similar effects on the spatial distribution of LELs, machine learning reveals these compounds can differ more subtly in how they alter LEL morphology.

**Fig. 4:**
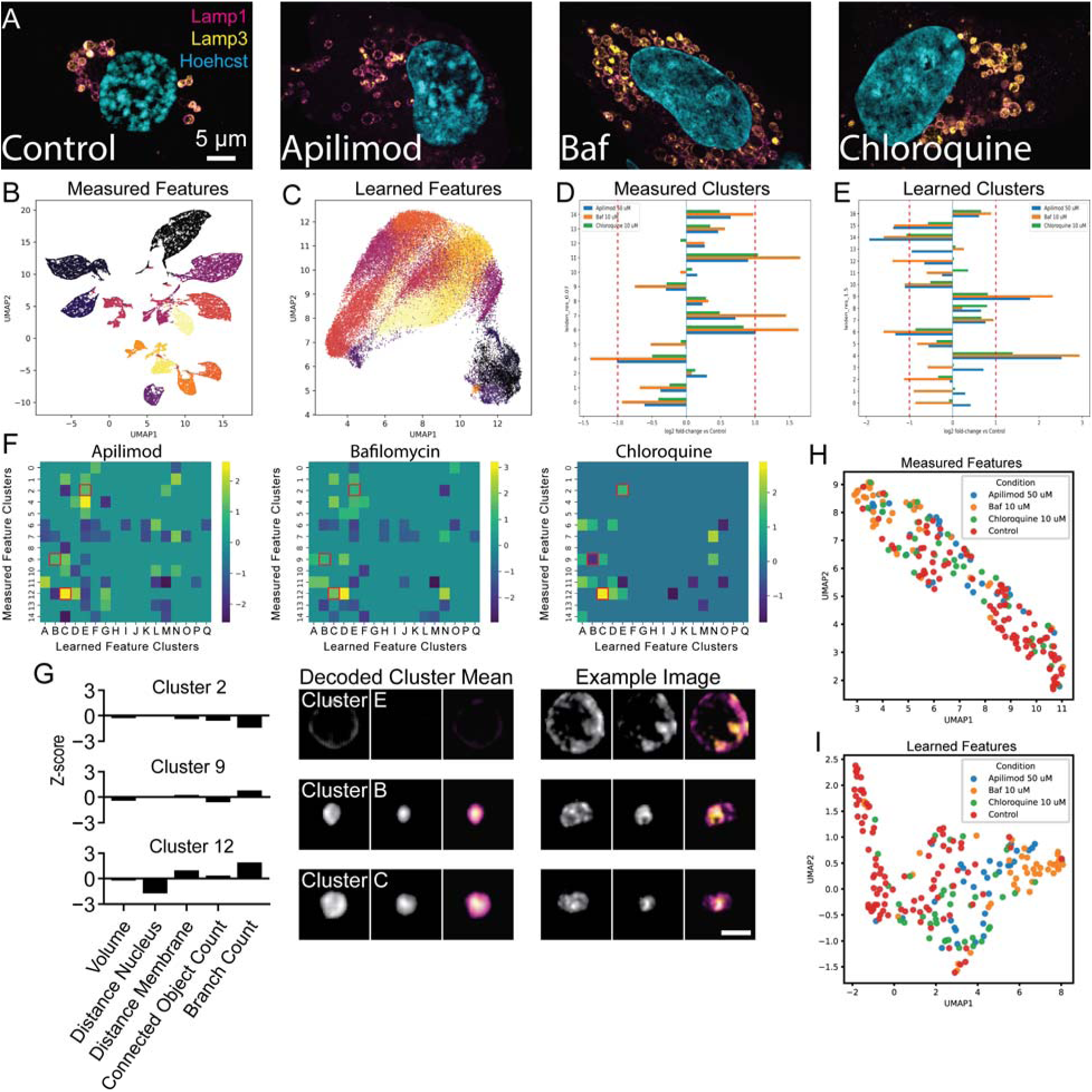
Comparison of Measured and Learned Features of LELs across Lysosomal Perturbations. A) Example whole-cell single-plane images from each of the noted perturbation conditions at their highest concentrations. B) UMAP of measured features across 73,890 LELs from all treatments in the perturbation panel (four replicate wells per treatment condition across two experiments with ∼10 fields-of-view per well capturing one cell each, 332 cells total). C) UMAP of learned features across LELs from all treatments in the perturbation panel. D) Log2 fold enrichment for occupancy of measured feature clusters compared to control for bafilomycin, apilimod, and chloroquine. Red dotted lines indicate a 2-fold change compared to control. E) Same as D, but for learned features. F) Delta-Log2 fold enrichment plots cluster comparison between measured and learned features in bafilomycin, apilimod and chloroquine compared with control. In each case, the enrichment/depletion table was generated as in Figure 3E, but for each of the perturbation conditions, then the same table from Figure 3E (representing the control condition) was subtracted from each. G) Representations of key cluster combinations from F. As in Fig. 3F, plots (left) identify measured feature clusters while image sets (right) represent learned feature clusters. Scale bar: 1 µm. H) Cell-level UMAP of measured features. To generate this plot, each cell was expressed as a vector, where each element is the proportion of its constituent LELs that belong to a given cluster in the measured feature data. I) Cell-level UMAP of learned features, generated the same way as H.

Having established a link between LEL spatial organization and morphology, we next asked whether these perturbations altered that relationship. To test this idea, we compared the changes in enrichment and depletion of measured-feature/learned-feature cluster pairings relative to control to identify perturbation-specific shifts in how morphological LEL states are distributed across spatial clusters (Fig. 4F). Apilimod, bafilomycin, and chloroquine share several enriched cluster combinations, consistent with the known effects of these compounds on lysosome function compared to control treatment with DMSO. Conversely, one cluster combination in particular is depleted in chloroquine-treated cells while being enriched in cells treated with either bafilomycin or apilimod (Fig. 4G). Thus, this cluster combination could distinguish the effects of apilimod and bafilomycin from those of chloroquine on LEL characteristics, suggesting that our approach could be used for scalable mapping of biologically relevant phenotypic perturbations at high resolution.

We next tested whether our individual LEL observations could be extrapolated to the single-cell level. Thus, we represented each cell by its distribution of LEL cluster identities (Fig. 4H,I). This analysis generates a compositional fingerprint of LEL state from the LEL population within each cell. Using this representation, learned-feature cluster occupancies produced greater separation between perturbation conditions than occupancies derived from measured-feature clusters, suggesting that the learned feature space captures treatment-specific LEL phenotypes at high resolution. Taken together, our work outlines a pipeline for HiExM that is ideally suited for mapping discrete features of individual organelles. This readily deployable approach could enable a deeper understanding of cell biology and can be integrated as an informative tool to aid in drug design and screening to improve therapeutic outcomes.

## Discussion

Cells can be understood in part through the properties and organization of their organelles. In this work, we developed a comprehensive experimental and analytical framework for analyzing individual organelles at scale, revealing quantitative information about morphology, spatial context, and perturbation response within and across cells. Looking forward, these methods could be extended beyond LELs to other organelles with structure–function relationships including mitochondria and nuclei. More broadly, the ability to quantify multiple organelle classes within the same cells may enable increasingly integrative representations of cellular state, in which cells are described by the coordinated behaviors of their organelles.

Microscopy-based cell biology has traditionally proceeded along two largely separate tracks: high-throughput studies that survey many conditions at relatively low spatial resolution, or lower-throughput studies that focus on a smaller number of cells in greater detail. Our HiExM pipeline closes the gap between these two tracks. We used LELs as a benchmark for this approach because of their known heterogeneity among cells. Importantly, LELs occupy a size range where the increase in resolution from HiExM can reveal morphological features at the single-organelle level. Additionally, these organelles are discrete and, for the most part, roughly spherical, making them amenable to single-organelle segmentation and formatting as uniform inputs for a machine learning model.

We demonstrate several advantages to our integrated approach. First, our quantitative pipeline provides information beyond conventional colocalization measurements. While colocalization is generally assessed based on broad signal overlap in standard confocal imaging^32,33^, our approach can better distinguish fluorescent signals on a per-organelle level at increased resolution beyond the diffraction limit of typical light microscopy. For example, we demonstrate that HiExM can overcome a limitation in therapeutic screening where methods for evaluating intracellular drug trafficking at scale often lack the spatial resolution needed to detect biologically meaningful differences in localization. Our results measuring differences in localization among three different anti-TfR antibodies within LELs over time suggest that incorporating higher-resolution, organelle-resolved trafficking analyses could improve the evaluation and optimization of intracellular delivery platforms. While much of our analysis used LAMP1 as a broad benchmark of LELs, an important future direction will be to further resolve how LEL functional identity maps to the learned feature space described in this work. Continued advances in universal staining procedures and optimized antibodies for ExM will be a critical next step.

Machine learning has transformed our ability to extract biological information from microscopy images. For example, the Allen Institute has published unsupervised models that learn reduced representations of whole cells from microscopy images^1,29^. Other studies have used machine learning to build classifiers for organelle state^34–37^. In this work, we extend these directions to the level of individual organelles by combining HiExM with unsupervised learning on segmented LELs. We use learned representations of individual LELs in our data to probe the relationship between organelle morphology and spatial positioning within the cell. Through this approach, we identified multiple spatially defined LEL states that are strongly associated with specific morphological classes in hCMECs. Further, when we perturbed lysosomes through three different mechanisms, we observed dramatic changes in the learned features that were masked in the measured features. In some cases, these changes differed between perturbations, suggesting that different perturbations can give rise to distinct LEL states even when they converge on a related lysosomal functional outcome. In the context of image-based drug screening, where viability or other narrow readouts are often used, our pipeline captures broader lysosome-associated changes in cell state. This information could benefit, as an example, cancer research, where lysosomes are increasingly implicated in therapeutic response and organelle-level phenotypes could reveal hidden effects that are missed by conventional screening readouts^38,39^. Together, the integrative framework developed in this work demonstrates how learned organelle features can be extracted from imaging data to reveal potentially hidden biological principles that can be broadly applied to understanding disease states and strengthening drug screening applications.

## Materials and Methods

### Cell Culture

Human cerebral microvascular endothelial cells (hCMEC/D3) were purchased from Coriell Institute (#CLU512) and cultured in accordance with the supplier’s protocols. Briefly, 96-well plates (CellVis #P96-1.5H-N) were coated with Collagen I (R&D Systems #3443-100-01) by incubating with a 1:25 dilution of Collagen I in DI water for 1 hour at 37°C. Cells were seeded at 15,000 cells per well and cultured for 4 days prior to experiments. Media changes were performed 24 hours and 72 hours after seeding.

### Immunofluorescence

For imaging, cells were fixed by first subjecting them to ice-cold MeOH for 5 minutes at −20°C. Next, MeOH was aspirated, ice-cold 1% PFA (Thermo Fisher Scientific #043368.9M, diluted in ice cold PBS) was added, and cells were left at 4°C for 30 minutes, followed by 2 PBS washes. Throughout the fixation process, care was taken to ensure that the plate remained ice-cold. The tandem MeOH/PFA fixation was used because MeOH fixation yields better Lamp1 staining than PFA alone, but MeOH alone is incompatible with expansion microscopy because MeOH does not confer any covalent linkages within the sample. Adding a light PFA fixation after the MeOH fixation step was compatible with expansion.

After fixation, cells were stored in PBS at 4°C until staining. For staining, cells were first blocked in a staining solution of 1% BSA (Millipore Sigma #A1595) and 0.1% saponin (Millipore Sigma #47036) in PBS for 5 minutes at room temperature. Following the blocking step, cells were incubated with primary antibodies (LAMP1: Cell Signaling Technologies #9091S; Lamp3: Developmental Studies Hybridoma Bank #H5C6; Cardiac Troponin T: Abcam #ab8295) in staining solution for 1 hour at room temperature, washed 3 times with PBS (one quick rinse, followed by a 5-minute wash, then a 10-minute wash), then secondary antibodies (Biotium #20125, #20105, and #20074) in staining solution for 1 hour at room temperature followed by the same washing regimen.

### Perturbation Experiments

For the compound perturbation experiments, hCMECs were subjected to chloroquine (MedChemExpress #HY-17589A), bafilomycin (Selleckchem #S1413), or apilimod (Selleckchem #S6414) at two different concentrations. For chloroquine and bafilomycin, cells were treated for 24 hours with either 100 nM or 10 uM. For apilimod, cells were treated for 1 hour with either 500 nM or 50 uM. These parameters were chosen based on previous studies using these compounds^30,31,39–45^. For cell volume identification, cells were transfected with an eGFP plasmid (Addgene #13031) using Opti-MEM and Lipofectamine 3000 (Thermo Fisher Scientific #31985062 and #L3000001, respectively) according to supplier protocols. Transfection was performed four days prior to the experiment.

### HiExM Protocol

Cells were subjected to expansion microscopy using the HiExM device and method as previously described with a few notable exceptions^10^. Briefly, cells were treated with 50 ug/mL Acryloyl X-SE (Thermo Fisher Scientific #A20770) diluted in ice cold PBS overnight at 4°C. The next morning, 100 uL of expansion gel solution was made from frozen stocks^40,41^. Specifically, 48 uL of 33% sodium acrylate (AK Scientific #R624), 24.5 uL of 25% 8-arm PEG Thiol (JenKem #A10022-1 / 8ARM(TP)-SH-10K), 3 uL of 25% crosslinker (Creative PEGWorks #PSB-3330), 7.5 uL of 40% Acrylamide (Millipore Sigma #A8887), 10 uL of 2% Lithium phenyl-2,4,6-trimethylbenzoylphosphinate (LAP, Millipore Sigma # 900889), and 7 uL of background solution (2M NaCl in PBS) were mixed together and vortexed. AcX solution was aspirated from cells and fresh expansion gel solution was applied to cells using HiExM devices (Supplementary Fig. 12). The gel solution was left on cells for 70 seconds, followed by irradiation with ultraviolet light (Uvitron International product # UV00002270, positioned 20 inches below the well plate and facing up) for 70 seconds. Finally, digestion solution containing 1 U/mL Proteinase K (New England Biolabs #P8107S) was added to the device-bearing well plate and left to stand for 6 hours at room temperature. Following a 6-hour digestion, Hoechst (Thermo Fisher Scientific #62249) diluted in DI water (1:2500) was added directly to wells containing digestion buffer and allowed to stand for 5 minutes. After this time, the whole well plate was submerged in ∼4L of DI water under constant stirring and allowed to stand overnight for expansion. The following day, excess water was aspirated from the wells of the well plate such that only the expansion gels remained. Next, 1.2% agarose (Genesee Scientific #20-102) was liquified in a microwave and then kept in a 37°C water bath to slowly cool down. Once the agarose was close to its sol-gel transition point, a transfer pipette was used to take agarose and then pipetted into the wells over top the expanded gels. Care was taken to ensure that the agarose would not flow (i.e. that it would not leave the pipet in drops but rather as a semi-gel continuous ‘snake’) so as to not allow any agarose to get underneath the expanded gel.

Once prepared, well plates were taken to an Opera Phenix (Perkin Elmer) confocal microscope. Samples post-expansion were imaged using the PreciScan protocol. For post-expansion, Hoechst was imaged with 60% power and 800 ms exposure time, and all other channels were imaged with 100% power 1000 ms exposure time. For pre-expansion, all channels were imaged at 50% power and 100 ms exposure time. All images were taken with 63x water immersion objective (NA 1.15) with 1 µm spacing between slices (note that this spacing does not constitute Nyquist sampling; we opted for under sampling in this work to increase image acquisition rate).

### Time-course Anti-TfR Antibody Experiments

For time-course experiments, cells were first incubated with serum-free media for 1 hour. After 1 hour, cells were incubated with media containing an anti-TfR receptor antibody at 180 nM as well as human transferrin at 30 nM(Millipore Sigma #T8158)^20^. Additions of serum-free media and antibody-loaded media were performed iteratively at 30-minute intervals for a total of 5 hours. After iterative loading of experimental treatments, all cells were fixed simultaneously and the plate was prepared for imaging.

### Image Analysis

All image analysis was performed in Python unless otherwise stated. All Python scripts used in this work can be found in the Boyer Lab GitHub at the following address: https://github.com/lboyerlab/Mapping-of-Organelle-Heterogeneity.

#### Segmentations

To segment individual LELs in images of expanded cells, we employed StarDist^15,16^. Notably, due to the variability in size of LELs in our datasets, we trained two separate models to identify LELs within different size ranges. Each model was trained on at least 8 ground-truth masks from manual segmentations generated in the Fiji plugin LabKit. These models were validated using code provided within the StarDist Jupyter Notebooks (Supplementary Fig. 2). For masking of individual cell volumes in Fig. 3, as well as masking of cTNT-positive cells in Supplementary Fig. 5, a simple adaptive threshold was employed based on cytosolic eGFP or cTNT signal, respectively.

#### LEL Metrics

The following describes how each of the five measured features of LELs were measured. To measure LEL volume, the number of voxels inside the segmented LEL mask was used. To measure the distance from the nucleus, the Hoechst-channel image was thresholded with an absolute threshold, then the distance from the segmented LEL to the nearest point on the binary mask of the nucleus was taken. To measure the distance from the membrane, first an adaptive threshold was applied to the eGFP stain, then the distance from the segmented LEL to the nearest background voxel in the binarized eGFP image was taken. For number of connected objects, the value simply represents the number of other LEL segmentations were immediately adjacent (ie. no background voxels separating them). Finally, for the number of branches in a minimal graph, first a minimal graph was constructed, where the nodes are the centroids of each LEL in 3D, and the edges were placed such that each node has at least one edge and the total edge length across the graph was minimized. Then, the number of edges connected to a given node was taken as the number of branches for the respective LEL.

#### Machine Learning Model Architecture and Training

We implemented a 3D β-variational autoencoder (β-VAE) in TensorFlow/Keras, following the general framework of Donovan-Maiye et al. (2022), adapted to StarDist-segmented two-channel fluorescence microscopy image crops of LAMP1-stained cells. The encoder comprised three 3D residual blocks with filter depths of [8, 16, 32], each consisting of a strided 4×4×4 convolution, a 3×3×3 convolution, batch normalization, and a residual skip connection via 1×1×1 pointwise convolution with average pooling. The flattened encoder output was projected to a 256-dimensional latent space via dual fully connected heads. The decoder mirrored the encoder using transposed convolutions with upsampling residual blocks, reconstructing the input volume through a final sigmoid-activated Conv3D layer with zero-padding and cropping to recover the original spatial dimensions. The KL divergence regularization loss was scaled by a β coefficient that was linearly annealed from 0 to a maximum value of β = 0.11 over training to prevent KL collapse — a phenomenon we observed in earlier model versions wherein the KL term suppresses the latent space before the encoder has learned a meaningful representation. The basic design of this model, as well as a full parameter set, can be found in the supplement (Supplementary Fig. 9, Supplementary Table 1).

#### Hyperparameter Optimization and Training Procedure

Hyperparameters were selected via Bayesian optimization using the Optuna library, which constructs a probabilistic model to guide the search toward promising regions of the hyperparameter space more efficiently than standard grid search methodology. The final configuration used a learning rate of 1.08×10^−4^, L2 regularization strength of 3.40×10^−5^, latent dimensionality of 256, and a batch size of 32. The model was compiled with the Adam optimizer (gradient clipping norm = 1.0) and mean squared error as the primary reconstruction loss; normalized MSE and structural similarity index (SSIM) were tracked as auxiliary metrics. The full dataset of 100,000 images was partitioned into training (∼80%), validation (∼10%), and test (∼10%) splits using stratified random splitting with a fixed random seed. Training proceeded for a maximum of 400 epochs with early stopping monitored on validation loss (patience = 20 epochs), restoring weights from the best-performing epoch. Experiment tracking, including hyperparameter logging and per-epoch metrics, was managed using MLflow. Model training was performed in Google Colab using an A100 GPU.

#### Object-Based Colocalization

To assess colocalization on a per-organelle basis, single organelle segmentations were taken from LAMP1 stains and applied to the drug channel of the image. Once the mask was applied, each individual LAMP1 volume with drug signal intensity data was saved as a tiff file, and the signal intensity information in those files was plotted.

## Supporting information

Table of Resources

## Acknowledgements

This work was funded by Sanofi iDEA iTech Award and the Open Innovation Lab (LAB), Deshpande Center (LAB), the KI Frontier Award (LAB), National Science Foundation Graduate Research Program Grant No. 2141064 (JHD), and the P30-CA014051 Koch Institute core support grant. Specifically, we thank the Koch Institute’s Robert A. Swanson (1969) Biotechnology Center High Throughput Sciences Core Facility (RRID:SCR_026340) and the Nanotechnology Core for scientific collaboration and expertise. We thank the Center for the Development of Therapeutics at the Broad Institute and acknowledge funding through the S10 grant NIH OD-026839–01. We thank Boyer lab members especially Victoria Tholkes, Abigail Lytton-Jean of the KI Nanotechnology Core and Andrew McCall (University of Buffalo) for helpful discussions. We thank the Anseth lab (University of Colorado, Boulder), especially Nathaniel Skillin, for help troubleshooting the PhotoExM gel. We are grateful to Sanofi team members including Karen Chandross, Mary Clare Mccorry, and Isaac Wolf for managing the collaboration as well as Julia Maeve Bonner and Susu Duan for intellectual contributions, technical recommendations, and material sharing.

## Data Availability

Data from this publication are available on Dryad. DOI: https://doi.org/10.5061/dryad.8gtht774b

## Author Contributions

Conceptualization: JHD, LAB. Methodology: JHD. Investigation: JHD, DY, FNN, GN, MA. Formal analysis: JHD, JF. Writing: JHD, LAB. Supervision: LAB. Funding acquisition: LAB. Resources: EA, AB, NCL, KWK.

