## Supplementary Information for "HiExM Enables Scalable Mapping of Organelle Morphology and Spatial Heterogeneity"

**CONTENTS:**

**FIGURES:**

Supplementary Figure 1: Non-Rigid Registration of LELs Show Typical Expansion Microscopy Metrics.

Supplementary Figure 2: Validation of StarDist Segmentation.

Supplementary Figure 3: Expansion Microscopy Resolves LEL Lumens.

Supplementary Figure 4: AbM-GAA Exhibits Greater LAMP1 Colocalization Compared to Cargo-Free Controls.

Supplementary Figure 5: HiExM Shows AbM-GAA Rescues Lysosome Hypertrophy and Sarcomere Structure in Pompe hiPSC-derived Cardiomyocytes.

Supplementary Figure 6: ToCLEM Identifies Morphologically Diverse LAMP1 Organelles.

Supplementary Figure 7: Measured Feature Cluster Identities by Feature Z-Scores.

Supplementary Figure 8: LEL Morphologies are Heterogenous within Cells.

Supplementary Figure 9: Variational Autoencoder Architecture and Validation Metrics.

Supplementary Figure 10: LAMP1 and LAMP3 signals show non-uniform distribution in learned feature space.

Supplementary Figure 11: Mean Vector Decodings for All Learned Feature Clusters.

Supplementary Figure 12: Photos of HiExM Gel Solution Collection System.

**TABLES:**

Table 1: Full Parameter Set for Autoencoder Model

Table 2: Compounds used for Lysosome Perturbations

The Table of Resources is provided separately as a standalone spreadsheet.

**SUPPLEMENTARY METHODS**

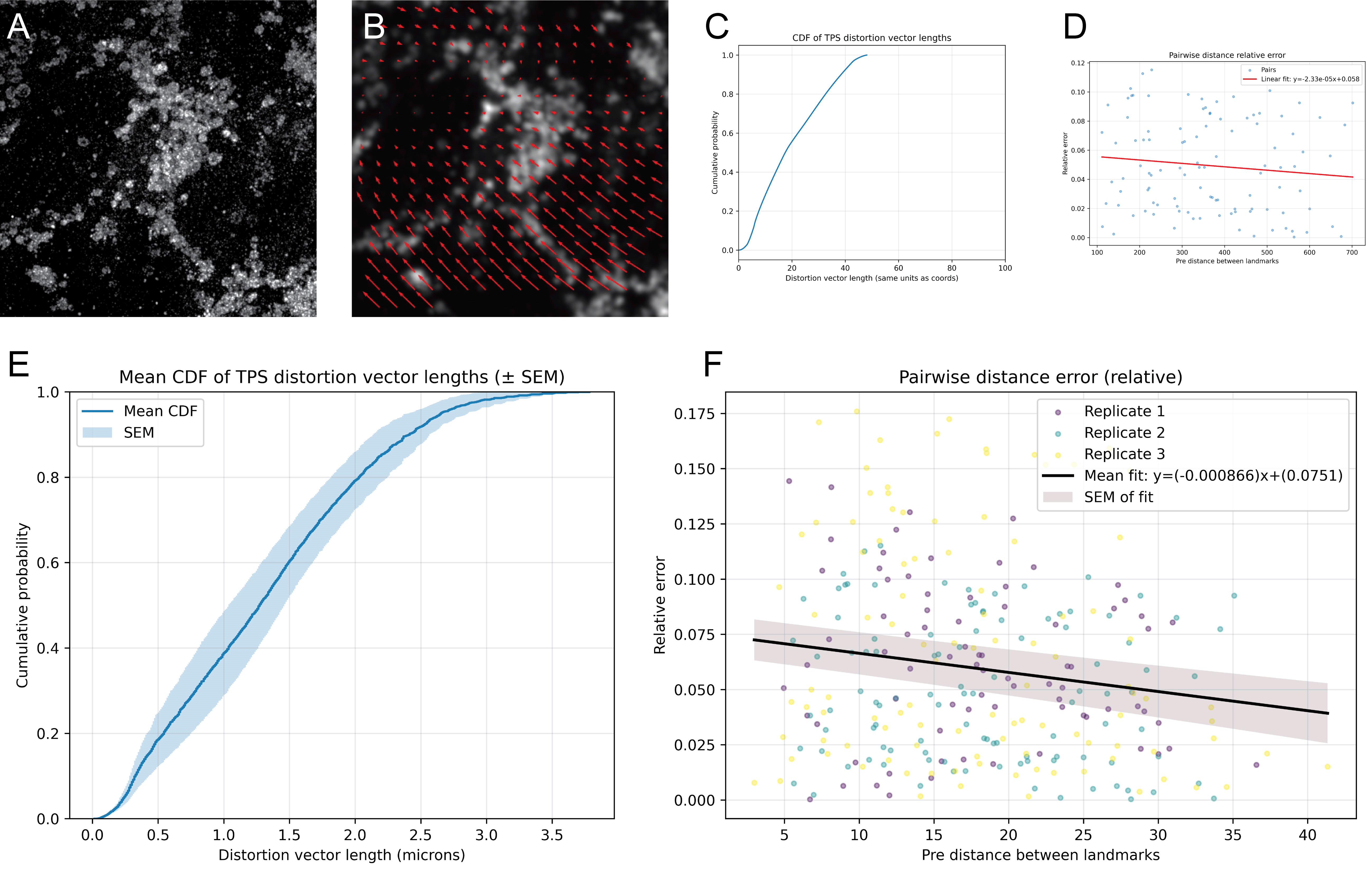
 ***Supplementary Figure 1: Non-Rigid Registration of LELs Show Typical Expansion Microscopy Metrics* (Related to Figure 1)*.*** A) Example post-expansion image of LAMP1. B) Corresponding pre-expansion image with overlayed vector-field mapping sample distortion. Thin-plate spline (TPS) deformation fields were estimated from corresponding landmark coordinates identified in pre- and post-expansion images and used to generate the vector fields. C) Cumulative distribution function (CDF) of vector lengths in the vector field. D) Relative pairwise distance error as a function of distance between landmarks. In this case, for each pair of landmarks the distance between those landmarks (pre-expansion) is expressed on the x axis and the relative error (measurement error post-expansion as a proportion of the measurement length) is expressed on the y axis. There is no significant relationship between the measured distance between two points and the relative measurement error. E) The CDF curves for vector length were measured for three images, and their mean curve was plotted as a shaded region representing standard error of the mean. F) Pairwise relative distance analysis was also repeated for three images and shows the same result as the example in D.

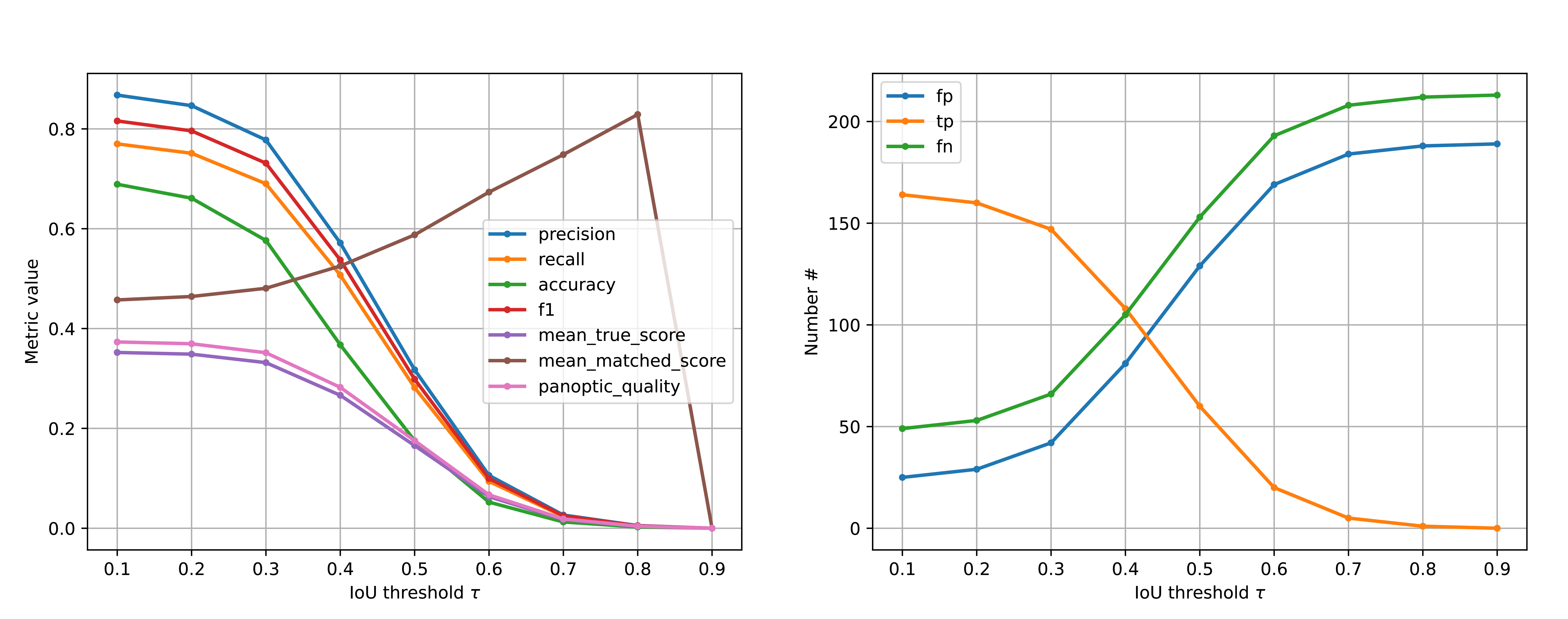
 ***Supplementary Figure 2: Validation of StarDist Segmentation*^1^ (Related to Figure 1)*.*** (Left panel) Several metrics of model quality are shown as a function of the intersection-over-union (IoU) threshold, which defines the minimum required overlap between a predicted object and a ground-truth object for that prediction to be considered a correct match. Precision indicates the fraction of predicted objects that correctly match a ground-truth object, such that higher precision reflects fewer false-positive detections. Recall indicates the fraction of ground-truth objects that are successfully detected by the model, such that higher recall reflects fewer missed objects. Accuracy summarizes the overall proportion of correct object assignments across predicted and ground-truth objects. The F1 score is the harmonic mean of precision and recall and therefore provides a balanced measure of detection performance when both false positives and false negatives are important. Mean true score represents the average overlap-based matching score across all ground-truth objects and therefore reflects how well the full population of true objects is recovered. Mean matched score represents the average overlap-based score only among successfully matched object pairs and therefore reflects the quality of segmentation for objects that were correctly identified. Panoptic quality provides a combined measure of detection and segmentation performance by incorporating both successful object identification and the quality of overlap between predicted and ground-truth masks.

(Right panel) The numbers of true positives (TP), false positives (FP), and false negatives (FN) are shown as a function of the IoU threshold, illustrating how the number of correctly identified objects, spurious detections, and missed objects changes as the matching criterion becomes more stringent.

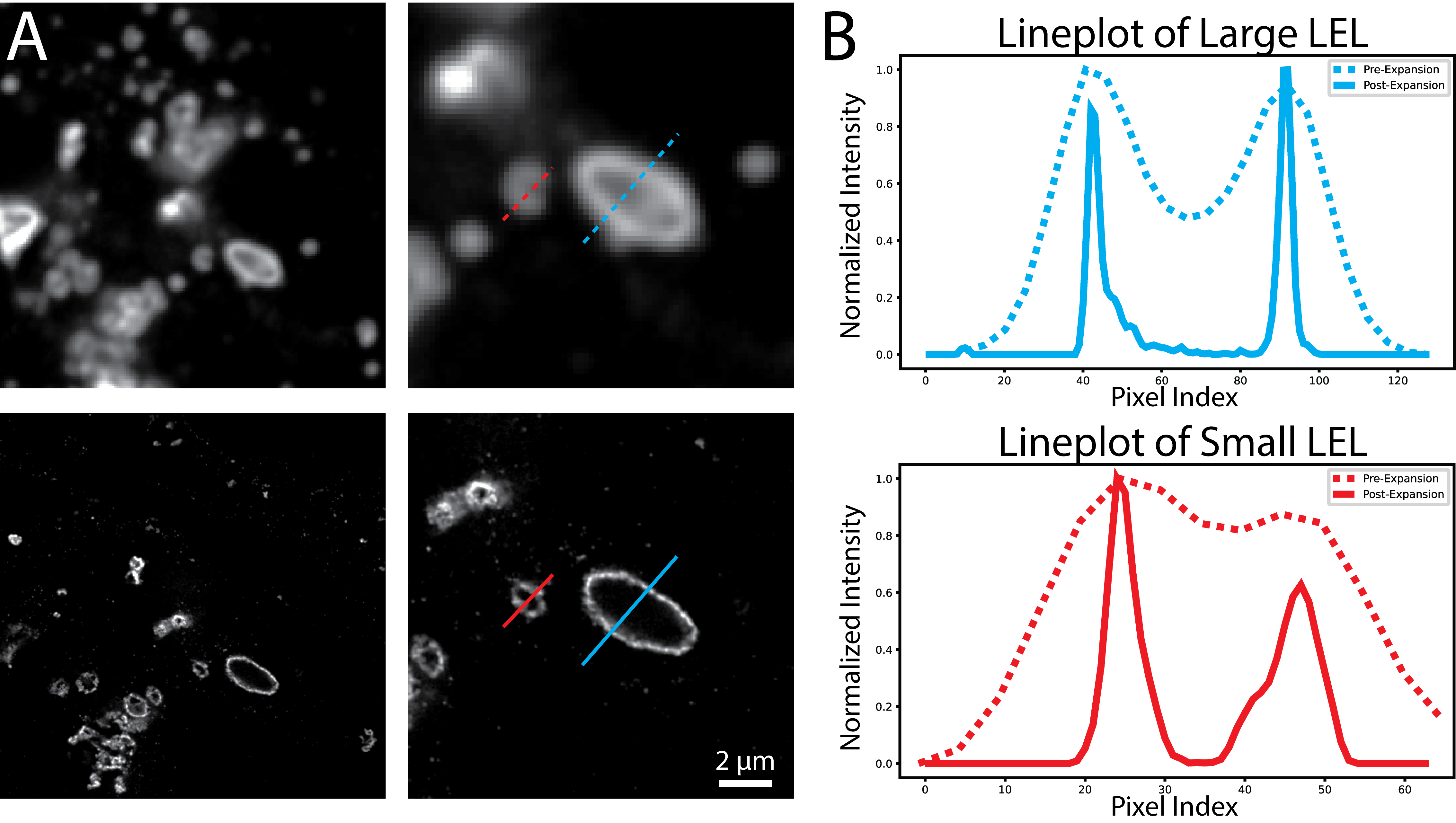

***Supplementary Figure 3: Expansion Microscopy Resolves LEL Lumens* (Related to Figure 1).** A) Example images of hypertrophic LAMP1 organelles from Pompe Cardiomyocytes that can present distinguishable lumens in standard confocal microscopy. Pre-expansion (top) and post-expansion (bottom) show that smaller organelles and clustered organelles are challenging to resolve with standard confocal microscopy. B) Line-plots across a small and large LEL showing the difference in resolving power between pre-expansion and post-expansion imaging.

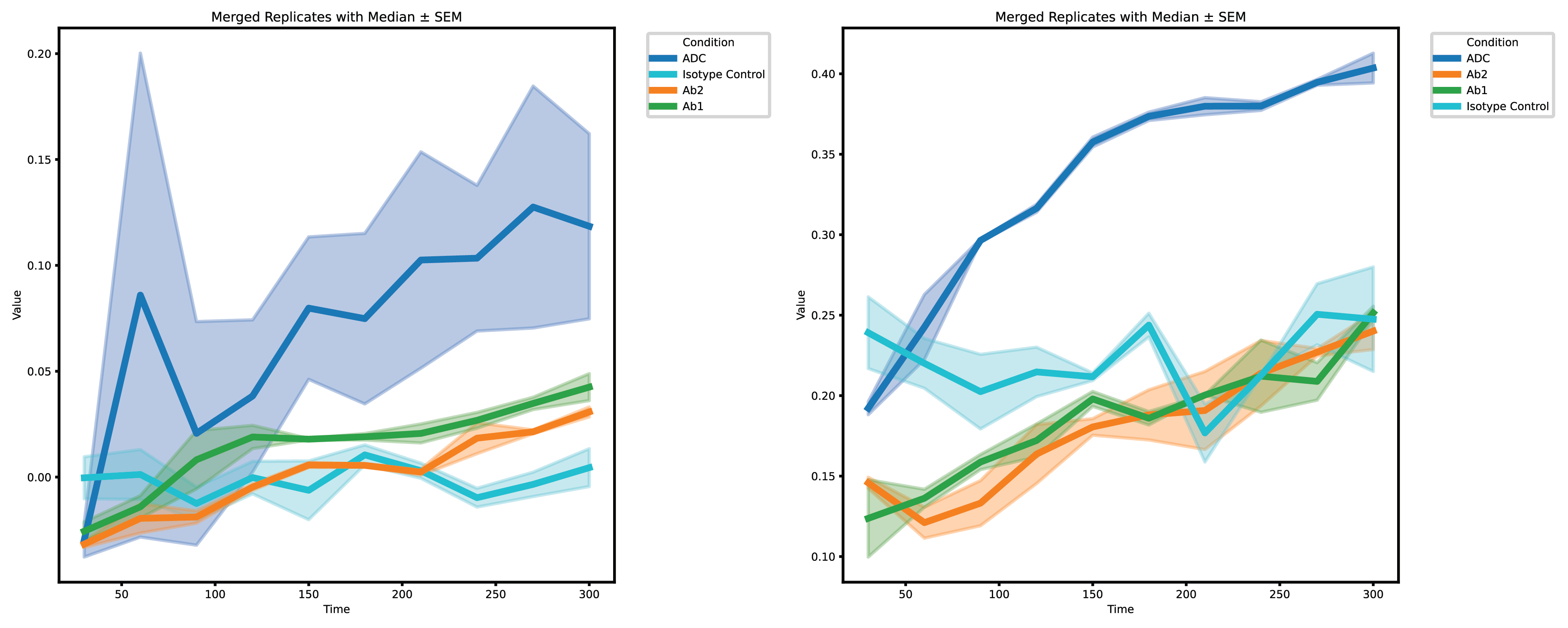
 ***Supplementary Figure 4: AbM-GAA Exhibits Greater LAMP1 Colocalization Compared to Cargo-Free Controls* (Related to Figure 2).** (Left Panel) Post expansion Pearson correlation coefficient over time for 4 different antibodies. (Right Panel) Pre expansion Pearson correlation coefficient over time for 4 different antibodies. Curves and shaded regions represent mean values and SEM values as in Figure 2. The isotype control is the same anti-TfR antibody as AbB but with no negligible binding affinity.

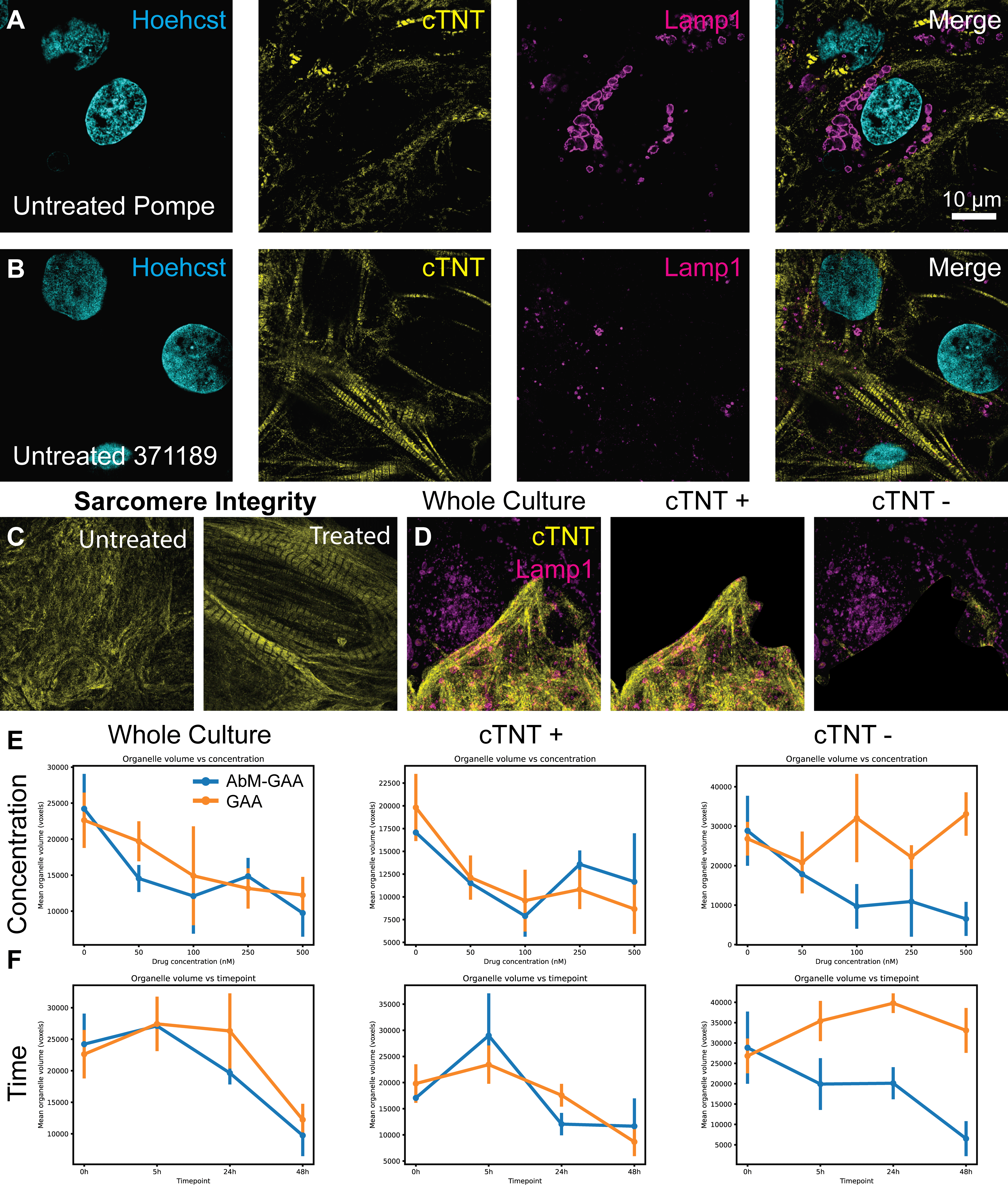

***Supplementary Figure 5: HiExM Shows ADC-GAA Rescues Lysosome Hypertrophy and Sarcomere Structure in Pompe hiPSC-derived Cardiomyocytes* (Related to Figure 2).** A) Example image (optical section) of untreated Pompe CMs. B) Example image (optical section) of untreated ‘Healthy’ CMs (371189 cells). C) Sarcomere structure is restored by treatment with both GAA and ADC. The image shown is an example from the ADC treatment and is representative of both conditions. D) Example image (max projection) of Pompe CMs showing the whole field of view (right), cardiac troponin T (cTNT) positive portion of the field of view (center) and the cTNT negative portion of the field of view (right). An adaptive threshold was used to separate the cTNT positive and cTNT negative areas. E,F) LEL volume is decreased by the ADC in a concentration-dependent (E) and a time-dependent (F) manner in cTNT negative cells. Plots showing the mean LEL organelle volume (voxel count) in whole fields of view (left), cTNT positive portions (center) and cTNT negative portions (right). All data is taken from the 48-hour timepoint of this experiment in E and from the 500 nM drug treatment in F. The stark difference between ADC and GAA in cTNT negative cells indicates that the TfR binding is necessary for the intended effect of GAA to manifest in these cells.

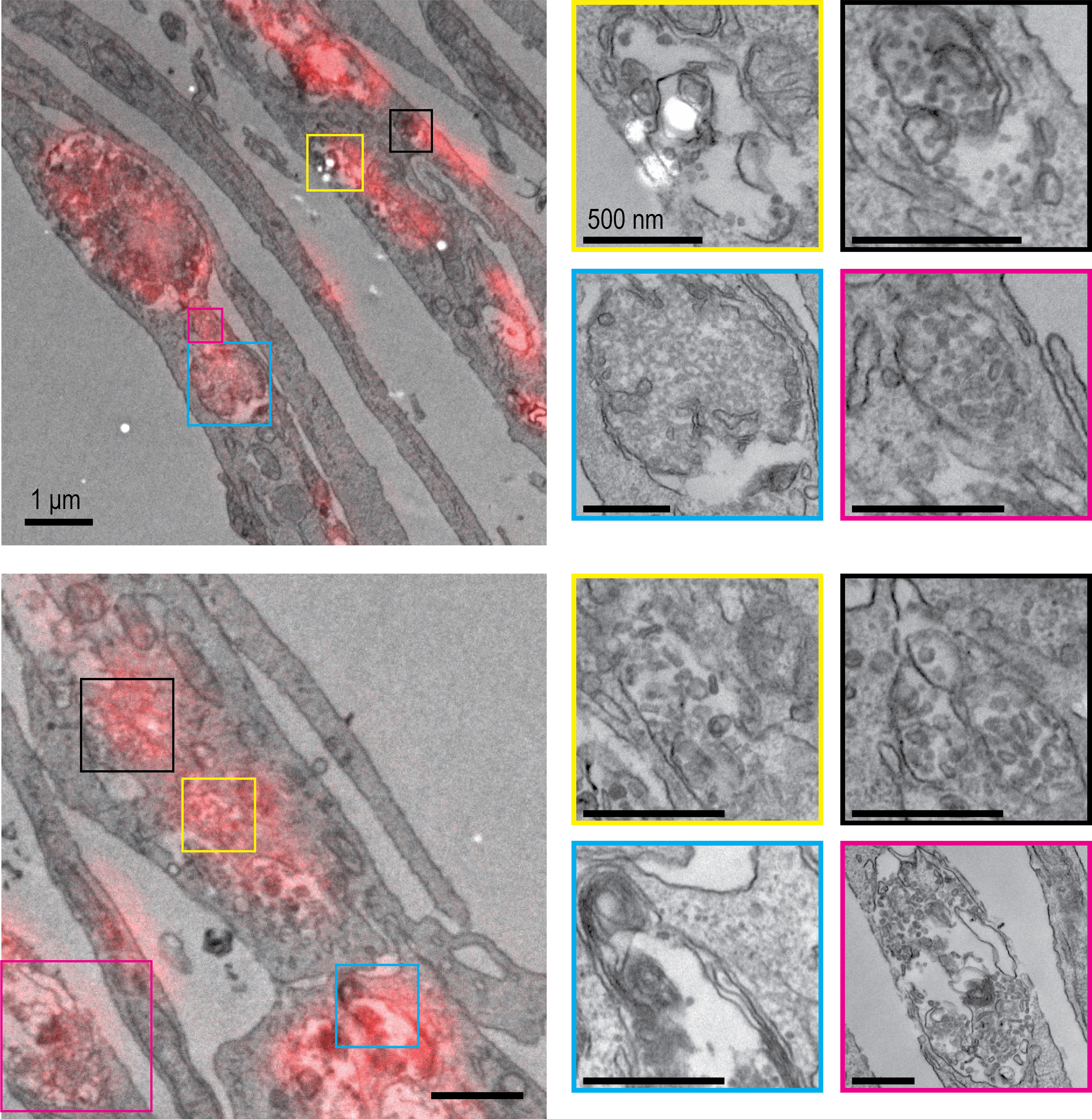
 ***Supplementary* *Figure 6: ToCLEM Identifies Morphologically Diverse LAMP1 Organelles* (Related to Figure 3).** (left) Two example images of hCMECs in TEM preparations with LAMP1 fluorescence stain overlayed. (right) High resolution insets from the corresponding CLEM images showing diverse presentation of LAMP1 positive organelles.

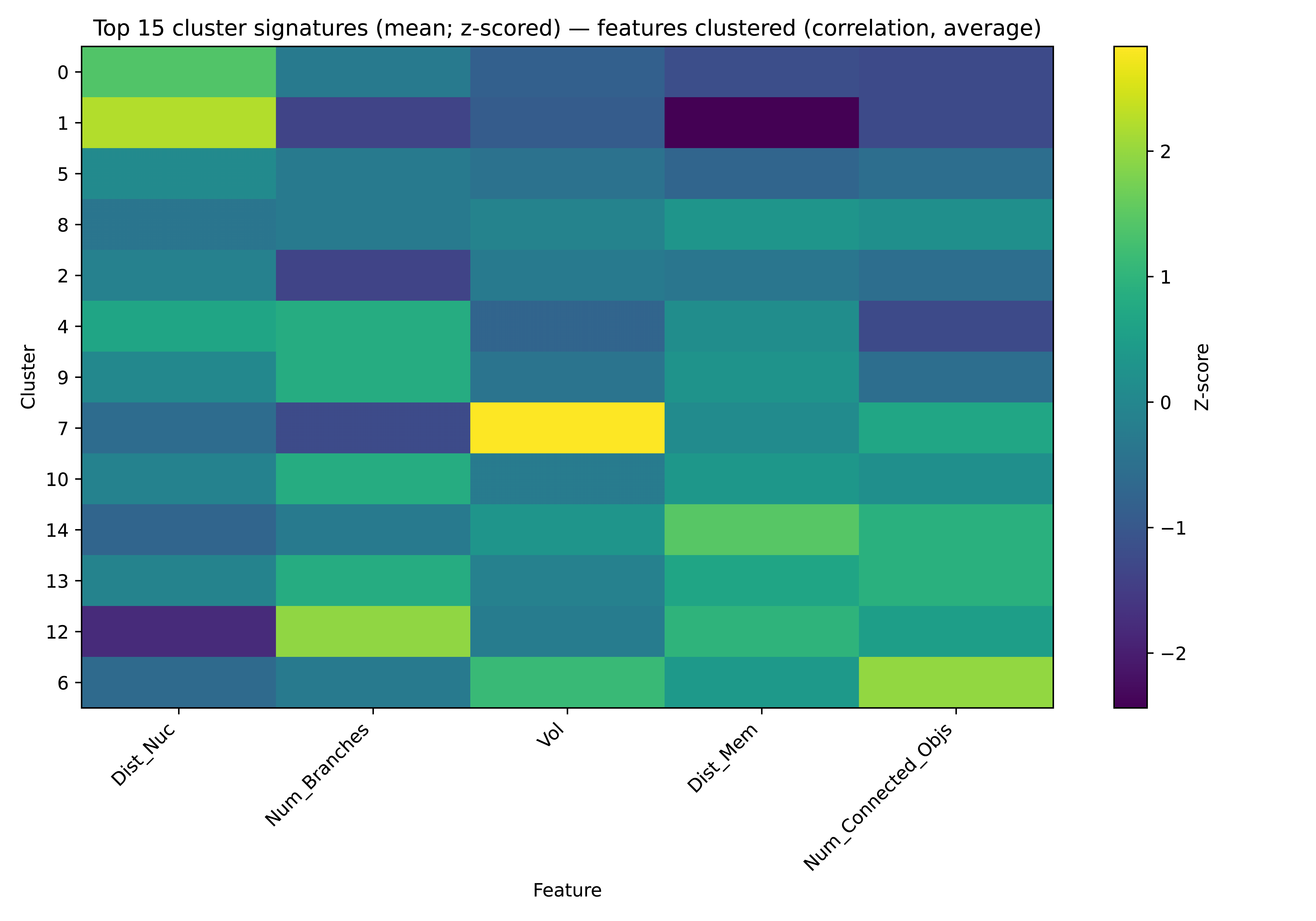

***Supplementary Figure 7: Measured Feature Cluster Identities by Feature Z-Scores* (Related to Figure 3).** Each row represents a cluster determined by Leiden clustering on the measured features from LELs. Those features include volume (voxel count), distance to the nucleus, distance to the membrane, number of connected objects, and number of branches in a minimal graph. The number of connected objects is equal to the number of unique label values in the StarDist mask that are touching the segmented organelle in question. The number of branches is calculated by generating a minimal graph, where the centroid of each LEL is a node. LELs with higher branch counts are more central in their local spatial environment with respect to other LELs, whereas higher connected object count indicates that the organelle is in a more spatially crowded environment.

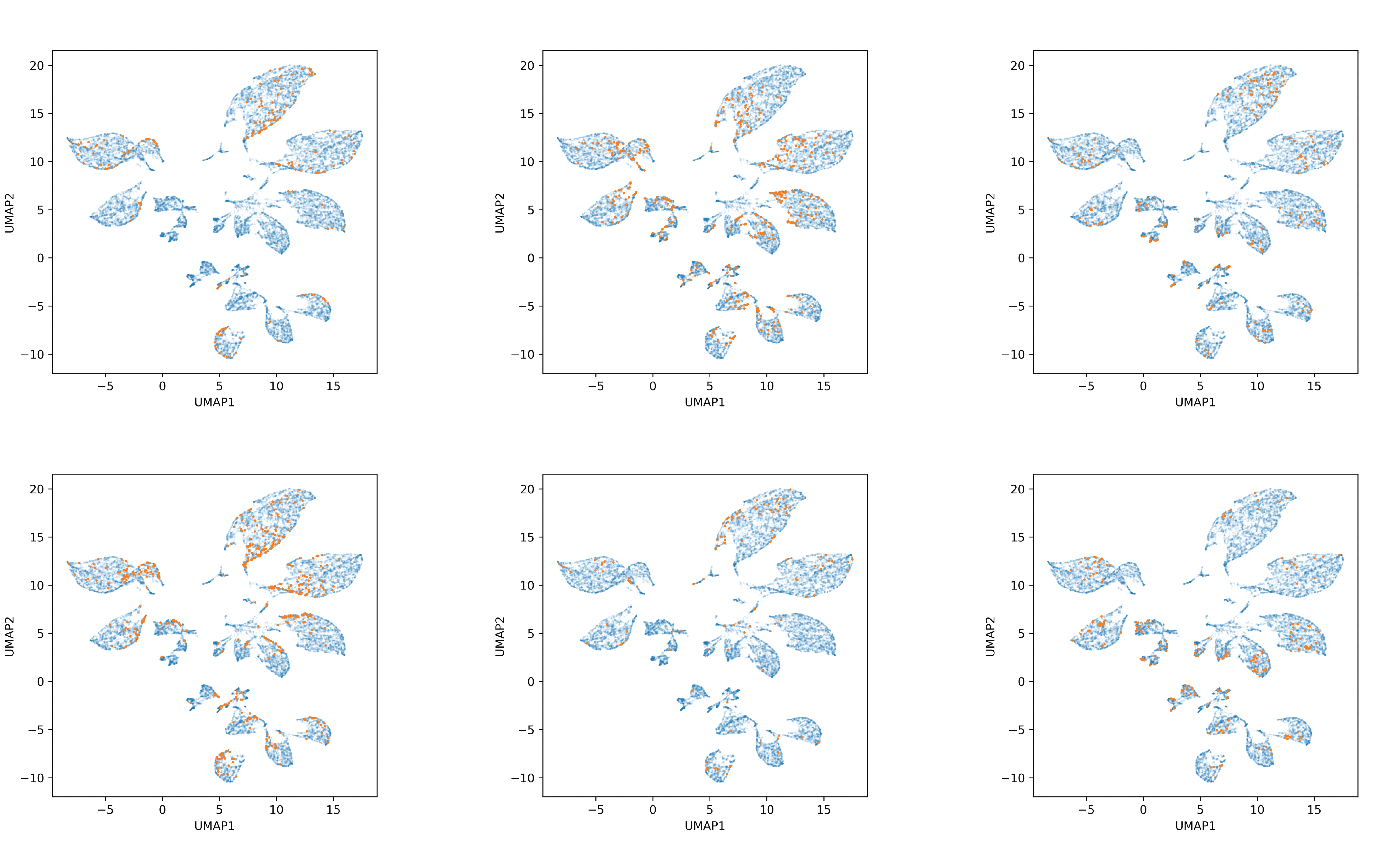
 ***Supplementary* *Figure 8: LEL Positioning is Heterogenous within Cells* (Related to Figure 3).** Six examples of single-cell hCMEC footprints (untreated control) by measured features, where each orange dot represents a single LEL for the given cell. In each case, the LELs distribute across the clusters shown in Figure 4B of the main text.

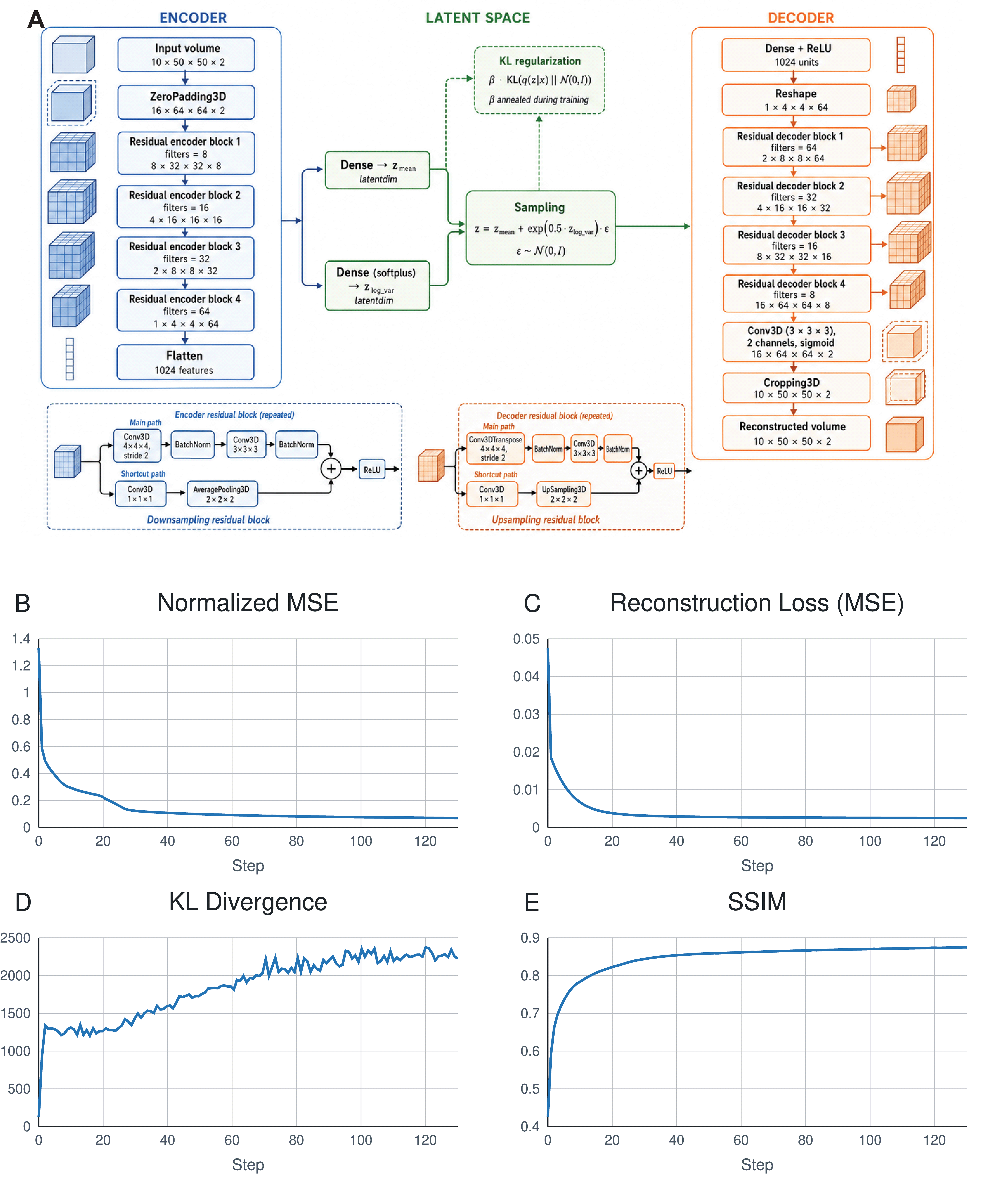

***Supplementary Figure 9: Variational Autoencoder Architecture and Validation Metrics* (Related to Figure 3).** A) Overview of model architecture. B) Normalized mean squared error (MSE) decreases smoothly and asymptotically, reflecting improvement in pixel-level reconstruction as the encoder and decoder refine their learned representations. C) Total reconstruction loss, comprising MSE and the 𝛽-weighted KL divergence penalty, also decreases smoothly and asymptotically. Early in the training cycle, the loss will closely track MSE (reconstruction loss) alone as 𝛽 is still held close to 0. As 𝛽 increases, the KL contribution starts to increase while the total loss continues its decline. This reflects the model learning to both minimize the reconstruction error while regularizing the latent space towards a standard normal distribution prior. D) KL divergence increases steeply during early epochs, indicating the encoder’s attempt to learn highly informative latent representations that deviate heavily from the standard normal prior while 𝛽 is still small. As 𝛽 increases, so too does the KL divergence, eventually reaching a plateau signifying the encoder has found an equilibrium between reconstruction quality and prior regularization. E) The structural similarity index measure (SSIM) exhibits the inverse behavior of the reconstruction and total loss curves. This is consistent with a rapid learning of the coarse structural features of our cell images followed by incremental refinement of finer details. Each plot shows per-epoch metrics computed on the validation set over the course of training.

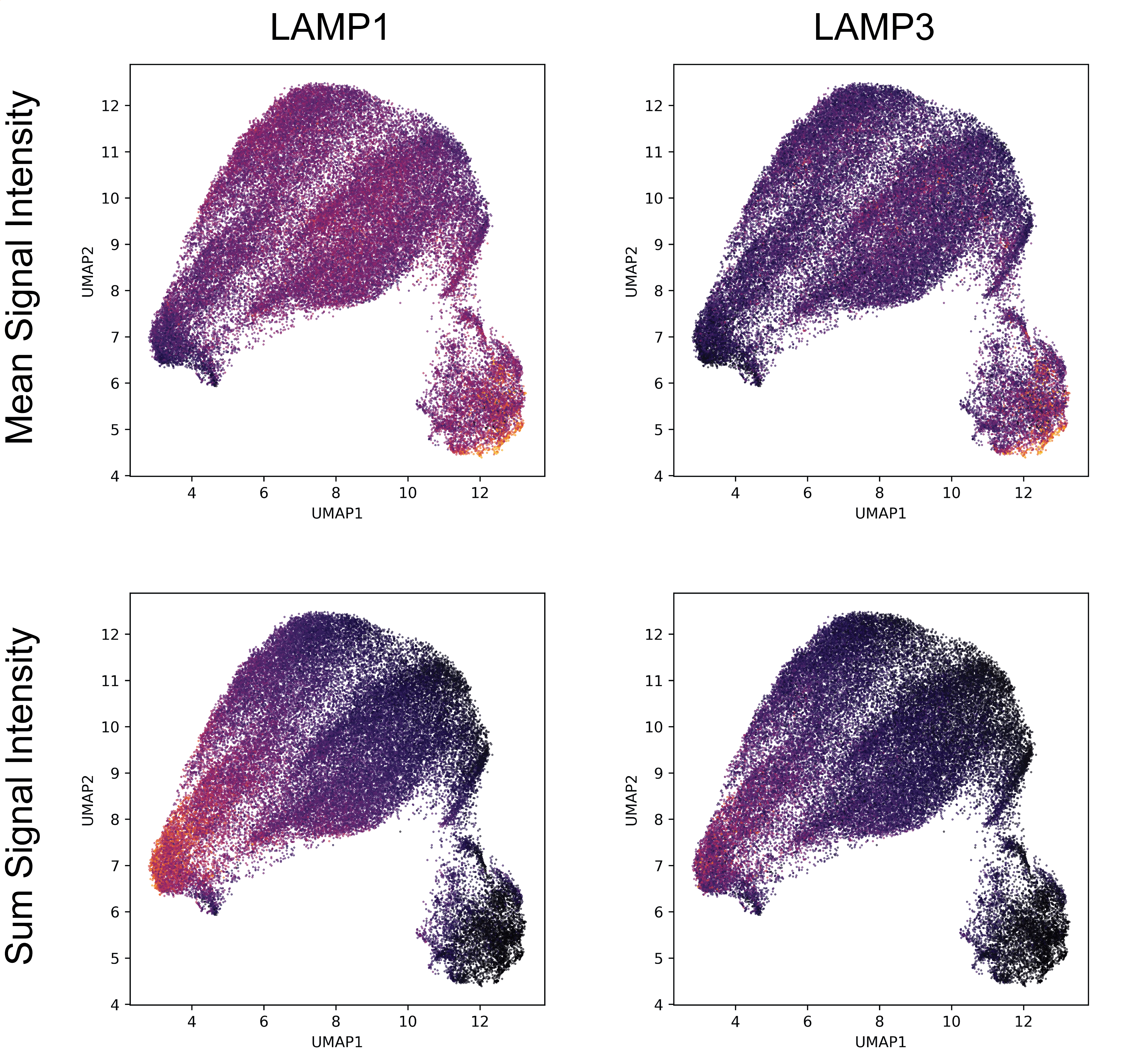

***Supplementary Figure 10: LAMP1 and LAMP3 signals show non-uniform distribution in learned feature space* (Related to Figure 3).** Four different representations of the learned feature space show the mean or sum signal intensity values of LAMP1 or LAMP3 for each organelle represented in the UMAP (each data point is a single organelle). Lighter color indicates higher value, darker color indicates lower value.

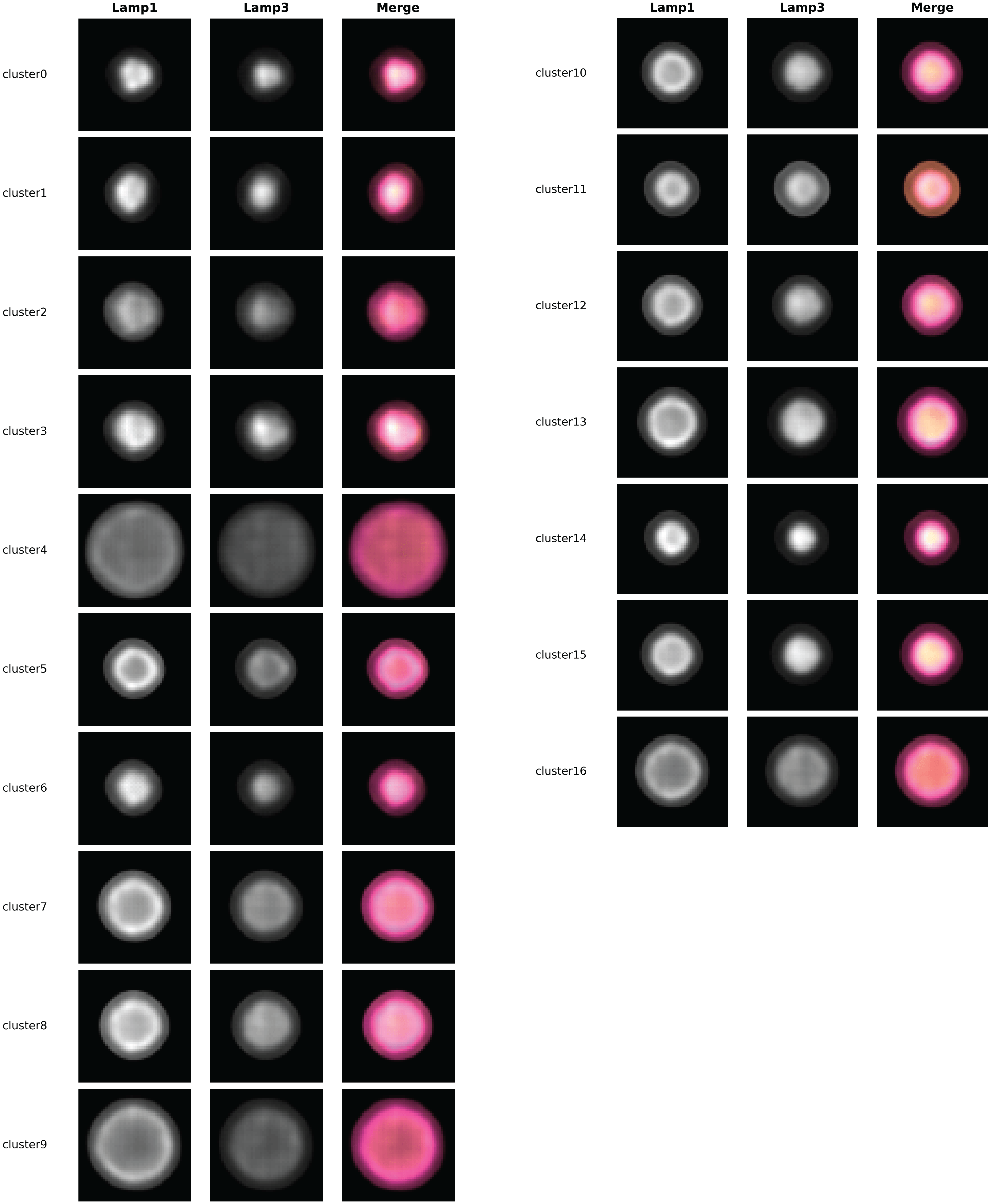

***Supplementary Figure 11: Mean Vector Decodings for All Learned Feature Clusters* (Related to Figure 3).** For each cluster in the learned feature space, the mean vector was taken for all of the data in that cluster, and the result was used to generate an image of the “average” organelle for that cluster using the decoding half of the variational autoencoder model.

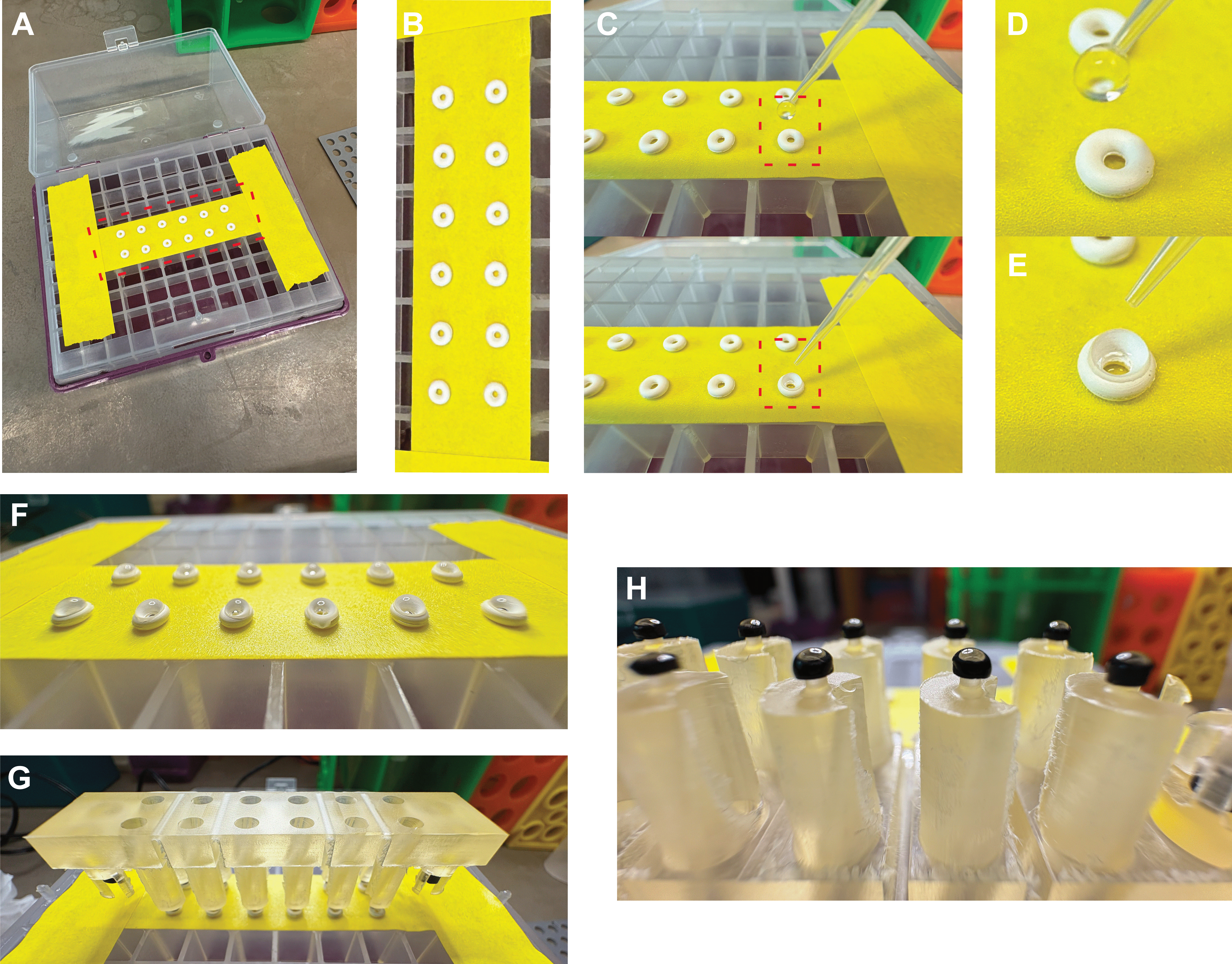

***Supplementary Figure 12: Photos of HiExM Gel Solution Collection System* (Related to Methods).** Part of the HiExM protocol was updated after the first publication to accommodate for the fact that PhotoExM gel is much more expensive per unit volume than the gel chemistry used in the initial work from the first paper^2–4^. Because of this, we modified the ‘gel reservoir’ to accommodate small volumes of PhotoExM gel. A) The gel reservoir now consists of a strip of upside-down lab tape secured tightly to the rack of a used pipette tip box. On the ‘sticky’ side of the tape (facing up), 12 o-ring silicone gaskets (McMaster-Carr #1284N102) are affixed by applying light downward pressure with a pair of tweezers. B) The resulting array of gaskets should be securely stuck to the tape surface and arranged in a 2 x 6 grid to match the grid of posts on the HiExM device. C) Next, the PhotoExM gel solution is pipetted onto the o-rings one at a time in 6 uL droplets. D) Gel solution is applied to the o-rings by dispensing the full 6 uL as a pendent droplet at the end of the pipet tip, then allowing the pendent droplet to make contact with the o-ring. E) The result is a dome shaped droplet resting centered on the o-ring. F) These droplets are dispensed onto all 12 o-rings. G) Next, the HiExM device is placed on the reservoir assembly such that each post meets one of the gel solution-loaded o-rings. H) When the device is removed, the posts retain small volumes of the gel solution. Note that the posts of the HiExM device now have black collars around their tips. These collars are made from heat shrink tubing cut into 1mm sections with a razor blade, and act to help the posts retain gel solution more consistently and effectively. After a device is loaded with gel solution, it is inserted into the well plate as previously described for the HiExM protocol, and the process is repeated with 5-6 more devices. 6 uL per o-ring is enough gel solution for 1 well plate worth of gels. If multiple well plates are prepared, the o-rings are reloaded with 4 uL of gel each between plates.

| Parameter | Value |
| --- | --- |
| Alpha | 0.8523464927212004 |
| Beta | 0.33384242768359496 |
| Batch Size | 32 |
| Input Dimension | (10, 50, 50, 2) |
| Latent Dimension | 256 |
| Learning Rate | 0.00010787701907600602 |
| Regularization Strength | 0.000033954412906648936 |

***Table 1: Full Parameter Set for Autoencoder Model.***

| **Feature** | **Bafilomycin (A1)** | **Chloroquine** | **Apilimod** |
| --- | --- | --- | --- |
| **Primary target** | Vacuolar-type H+-ATPase (V-ATPase) | Lysosomes (weak base; multiple targets) | PIKfyve (phosphatidylinositol-3-phosphate 5-kinase) |
| **Mechanism of action** | Direct inhibitor of V-ATPase → blocks proton pumping into lysosomes/endosomes | Accumulates in acidic organelles → buffers pH (raises lysosomal/endosomal pH) | Inhibits PIKfyve → disrupts PI(3,5)P₂ synthesis → impairs endolysosomal trafficking/maturation |
| **Effect on lysosomal pH** | Strong increase (prevents acidification) | Moderate increase (neutralizes acidic compartments) | Indirect; may alter lysosomal function but not primarily by pH neutralization |
| **Effect on autophagy** | Blocks autophagosome–lysosome fusion/function (late-stage autophagy inhibition) | Inhibits autophagic degradation by raising lysosomal pH | Disrupts endolysosomal trafficking; can indirectly impair autophagy flux |
| **Effect on endosomes/lysosomes** | Prevents acidification and maturation | Causes swelling and dysfunction of lysosomes | Causes enlarged vacuoles; blocks endosome-to-lysosome trafficking |
| **Specificity** | Relatively specific V-ATPase inhibitor | Less specific; pleiotropic effects | More selective molecular target (PIKfyve) |
| **Typical research use** | Gold standard for inhibiting lysosomal acidification | Widely used lysosomal/autophagy inhibitor; also clinical drug | Tool compound for studying endolysosomal trafficking and viral entry |
| **Clinical use** | No (too toxic) | Yes (antimalarial, autoimmune diseases) | Investigational/clinical trials (e.g., immune and viral contexts) |

***Table 2: Compounds used for Lysosome Perturbations.***

***Supplementary Methods:***

*Correlative Light-Electron Microscopy*

*Fixation and Sample Preparation.* All reagents were prepared fresh on the day of the experiment. Cells were initially fixed by adding an equal volume of 4% paraformaldehyde (PFA; Electron Microscopy Sciences, 15700) directly to the culture media, resulting in a final PFA concentration of 2%. This mild fixation was carried out for 30 min at room temperature. The fixation solution was then replaced with 4% PFA in 0.1 M PIPES buffer and samples were further fixed for 24 h at 4°C. Following fixation, cells were washed three times with 0.1 M PIPES (Sigma, P6757) and quenched for 10 min in 0.15% glycine (Sigma, G8790). Cells were subsequently harvested by scraping, pelleted by gentle centrifugation (600 × g, 5 min), infiltrated with 12% gelatin (Sigma, G2500) for 30 min at 37°C, and solidified on ice for at least 30 min. Gelatin-embedded cell pellets were cut into small blocks (~0.5 mm³) and infiltrated overnight at 4°C in 2.3 M sucrose (Sigma, S0389) under gentle agitation. The blocks were then mounted on cryo-ultramicrotomy specimen pins (Electron Microscopy Sciences, 75959-06), frozen in liquid nitrogen, and stored until sectioning

*Cryo-Ultramicrotomy.* Sectioning was performed using a Leica UC7 cryo-ultramicrotome equipped with a cryo-diamond knife (Diatome, cryo-immuno 35°). 1 µm sections were cut at -80°C and collected onto 12 mm glass coverslips using a mixture of equal parts of 2.3 M sucrose and 2% methylcellulose (Sigma, M6385). Sections were stored at room temperature until further processing.

*Fluorescence Imaging.* For immunofluorescence, coverslips with cryosections were incubated in 0.1 M PIPES at 37°C for 30 minutes to remove the sucrose-methylcellulose layer. Samples were then blocked in 1% BSA-c (Aurion, 900.099) in 0.1M PIPES for 30 min at RT before being incubated with a conjugated primary antibody against (add here antibody information). Finally, coverslips were washed three times in 0.1 M PIPES, followed by a final wash in deionized water (ddH₂O). and mounted on a glass slide using DAPI-containing mounting media (Invitrogen, D3571) for nuclear staining. The samples were imaged at a Zeiss LSM 880 confocal microscope.

*Electron Microscopy and Image Correlation.* After fluorescent imaging, the slides were unmounted and stained with heavy metal for EM. Briefly, the samples were exposed to reduced osmium (1% osmium tetroxide, 1.25% potassium ferrocyanide in 0.1M PIPES) for 15 minutes, washed three times in 0.1M PIPES buffer, and then stained again in 1% TCH (Thiocarbohydrazide) (Sigma, 223220) for 15 minutes. The sample was then washed three times in water before being stained again in 1% osmium tetroxide for 15 minutes on ice, and after 3 washes in water again in 1% uranyl acetate in 50 mM maleate buffer pH 5.5. After a final wash in water the sample was dehydrated in increasing ethanol concentrations, and resin infiltration (Embed 812 embedding kit with BDMA, EMS# 14121). All staining and dehydration steps were performed on ice. The infiltrated samples were baked for at least 48 hours at 60 C before being trimmed and sectioned with a Leica UC7 ultramicrotome. A diamond knife (Diatome) was used to cut ultrathin sections (50 nm), which were then collected in carbon coated formvar slot grid (EMS, FCF2010-Cu-EA), and imaged in a Technai T12 operating at 80 KV. The LM and EM images were then correlated using the Fiji plugin Big Warp

*hiPSC Culture and Cardiomyocyte Differentiation*

PENN0141-37-3 (37-1189) hiPSC line was generously provided by the University of Pennsylvania under an MTA agreement and was used as the wild-type control in this study. The lysosomal storage disease hiPSC line TRNDi007-B (also known as HT521B) was obtained from the Coriell Institute for Medical Research and is derived from skin fibroblasts of a five-month-old male infant with infantile-onset Pompe disease (glycogen storage disease type II), carrying a homozygous **c.2560C>T (p.R854X) mutation in the GAA gene, which results in deficiency of lysosomal acid α-glucosidase and intracellular glycogen accumulation characteristic of this lysosomal storage disorder^5^.

hiPSCs were maintained on plates coated with Matrigel (Corning, Cat. #354277) in mTeSR™ Plus media (STEMCELL Technologies, Cat. #100-0276) at 37 °C in a humidified atmosphere containing 5% CO₂, with media changes every two days. Cells were passaged using ReLeSR™ (STEMCELL Technologies, Cat. #100-0483) according to the manufacturer’s protocol.

Differentiation of hiPSCs into cardiomyocytes (hiPSC-CMs) was performed as previously described^6^. Briefly, cells were dissociated using Accutase (Innovative Cell Technologies, Cat#. AT104500) and seeded in a 12-well plate coated with Matrigel at a density of 1×10^5^ cells per well in mTeSR Plus media supplemented with 10 µM ROCK inhibitor (Millipore Sigma, Cat. #5092280001). The following day, ROCK inhibitor was removed and daily media changes were performed to allow cells to grow for 2-4 days until reaching 80-90% confluency. At day 0, cells were treated with CHIR-99021 (12 µM, Selleck Chemicals, Cat. #S2924) for 24 h in RPMI 1640 media (Thermo Fisher Scientific, Cat. #11875093) supplemented with B-27 minus insulin (Thermo Fisher Scientific, Cat. #A1895601). At day 3, half of the media was replaced with fresh media, resulting in a final concentration of 5 µM Wnt inhibitor IWP 2 (Tocris Bioscience, Cat. #3533). Forty-eight hours later, IWP2 was removed, and at day 7, cells were switched to RPMI 1640 media supplemented with B27 (Thermo Fisher Scientific, Cat. #17504044). At day 14, cardiomyocytes were dissociated using the STEMdiff™ Cardiomyocyte Dissociation Kit (STEMCELL Technologies, Cat. #05025), replated onto Matrigel-coated plates, and maintained for downstream studies.

*Quantitative Analysis of LELs in Pompe Cardiomyocyte Cultures*

To evaluate phenotypic rescue of lysosomal hypertrophy in Pompe CMs, we dosed CMs at day 18 with a range of concentrations (0, 50, 100, 250, 500 nM) of either enzyme alone or ADC and fixed after 6, 24, and 48 hours. To analyze LEL volumes, we used StarDist as previously described to segment individual LEL organelle volumes; as with hCMECs, we found that LELs were only able to be robustly segmented in post-expansion samples. We compared the distribution of LEL volumes between enzyme and ADC treatments as functions of concentration and time.

We stained for cardiac troponin T (cTNT) to separate cardiomyocytes from non-cardiomyocyte cells in our image data and compared LEL volumes from our experiments in both populations of cells. To separate segmented organelles in cTNT+ and cTNT- regions, we used an adaptive threshold on the cTNT channel to generate a binary mask of the generally cTNT volume constituting a reasonable approximation of the cytosolic volume for cardiomyocytes in the culture. Next, we collected the LEL segmentations overlapping with the cTNT mask and compared their volumes with the segmentations that were outside of the mask.
